# CASC3 promotes transcriptome-wide activation of nonsense-mediated decay by the exon junction complex

**DOI:** 10.1101/811018

**Authors:** Jennifer V. Gerbracht, Volker Boehm, Thiago Britto-Borges, Sebastian Kallabis, Janica L. Wiederstein, Simona Ciriello, Dominik U. Aschemeier, Marcus Krüger, Christian K. Frese, Janine Altmüller, Christoph Dieterich, Niels H. Gehring

## Abstract

The exon junction complex (EJC) is an essential constituent and regulator of spliced messenger ribonucleoprotein particles (mRNPs) in metazoans. As a core component of the EJC, CASC3 was described to be pivotal for EJC-dependent nuclear and cytoplasmic processes. However, recent evidence suggests that CASC3 functions differently from other EJC core proteins. Here, we have established human CASC3 knockout cell lines to elucidate the cellular role of CASC3. In the knockout cells, overall EJC composition and EJC-dependent splicing are unchanged. A transcriptome-wide analysis reveals that hundreds of mRNA isoforms targeted by nonsense-mediated decay (NMD) are upregulated. Mechanistically, recruiting CASC3 to reporter mRNAs by direct tethering or via binding to the EJC stimulates mRNA decay and endonucleolytic cleavage at the termination codon. Building on existing EJC-NMD models, we propose that CASC3 equips the EJC with the ability to communicate with the NMD machinery in the cytoplasm. Collectively, our results characterize CASC3 as a peripheral EJC protein that tailors the transcriptome by promoting the degradation of EJC-dependent NMD substrates.

## Introduction

Messenger RNA-binding proteins (mRBPs) determine the stability, location, efficiency of translation and fate of bound mRNAs and are therefore important regulators of post-transcriptional gene expression (1). A central component of spliced mRNPs in metazoans is the exon-junction-complex (EJC), which is deposited during splicing upstream of exon-exon boundaries (2–4). The heterotetrameric core of the EJC is composed of the proteins EIF4A3, MAGOH, RBM8A (Y14) and CASC3 (BTZ, MLN51) (5, 6). Generally, EJCs serve on spliced mRNAs as a mark that act as a binding platform for peripheral EJC-interacting factors (7). The core and peripheral EJC components contribute to different steps of post-transcriptional gene expression including splicing regulation, mRNA localization, translation and nonsense-mediated mRNA decay (NMD) (2, 4).

The EJC does not form spontaneously, but instead undergoes stepwise assembly in association with the spliceosome, while it proceeds through different spliceosomal complexes (8). As a first step, the splicing factor CWC22 recruits EIF4A3, to which the MAGOH/RBM8A heterodimer binds later on (9–13). Unlike the three spliceosome-associated EJC components EIF4A3, MAGOH and RBM8A, the fourth protein CASC3 is not detected in the purified spliceosomal C complex (14, 15). Furthermore, it cannot be detected in mRNPs formed on splicing intermediates (12). Therefore, it was suggested that CASC3 binds to the initially formed trimeric pre-EJC (consisting of EIF4A3, MAGOH and RBM8A) at a later stage. Interestingly, CASC3 has also been shown to be a shuttling protein that is mainly located in the cytoplasm, whereas the other EJC components are predominantly detected in the nucleus (16–20). It has therefore been suggested that CASC3 binds to the EJC in the nucleus and is transported with it into the cytoplasm (21). A recent study demonstrated that EJCs undergo a compositional switch and that the ASAP/PSAP component RNPS1 and the protein CASC3 bind to functionally different mRNPs and exist in mutually exclusive EJCs (22). While EJCs of nuclear enriched transcripts were found to interact with RNPS1, the EJCs of cytoplasmic enriched transcripts rather contained CASC3. This observation is in line with the predominantly cytoplasmic localization of CASC3, but would argue against a nuclear function.

Another aspect under debate is the involvement of CASC3 in the NMD pathway. According to the EJC-dependent model of NMD, an EJC present more than 50-55 nucleotides downstream of a premature termination codon (PTC) triggers degradation by the NMD machinery (23). This quality control mechanism rids the cells of aberrant transcripts that contain PTCs due to mutations or mis-splicing. Additionally, it serves as a post-transcriptional mechanism of gene expression, especially when coupled to alternative splicing (24). This is a common feature of many genes coding for mRBPs, e.g. most SR proteins (25, 26). The EJC triggers NMD by interacting with members of the SURF complex resulting in phosphorylation of the central NMD factor UPF1 (27). Phosphorylated UPF1 then stimulates two distinct degradation pathways of NMD: The SMG5/7-dependent pathway results in deadenylation and decapping of the transcript followed by exonucleolytic decay from the 5′ end by XRN1 and the 3′ end by the exosome (28). Alternatively, the transcript can be cleaved in the vicinity of the PTC by the endonuclease SMG6 which results in two mRNA fragments that can be exonucleolytically degraded by XRN1 and the exosome (29, 30). While these pathways can act redundantly and in principle compensate for each other, SMG6-dependent endonucleolytic cleavage (endocleavage) has been shown to be the dominant pathways for NMD in human cells (31–33). In cells depleted of CASC3 a stabilizing effect on PTC-containing reporter mRNAs and selected endogenous targets was reported (12,17,34). Furthermore, tethering CASC3 to an mRNA results in UPF1-dependent degradation of the transcript (12). However, a recent report has challenged these observations and showed that CASC3 plays a minor role in NMD and only for certain endogenous targets in contrast to EIF4A3 or RNPS1 (22).

We were intrigued by the contrasting reports about the enigmatic role of CASC3 and decided to investigate the function of CASC3 and its distinction to the other EJC core components in more detail. For this purpose, we established HEK 293 CASC3 knockout (KO) cells using CRISPR-Cas9-mediated gene editing. The CASC3 KO cell lines are largely unchanged in their composition of the EJC core and peripheral interacting proteins. However, RNA-sequencing reveals an upregulation of transcript variants containing premature-termination codons (PTC) as well as the differential expression of many known NMD targets, indicating a perturbation of this decay pathway. Mechanistically, CASC3 stimulates SMG6-dependent turnover of NMD targets and likely acts as a link from the EJC to the NMD machinery. On the basis of these results we propose a revised model of EJC-dependent NMD in human cells.

## Materials and Methods

### Cell culture

Flp-In 293 T-REx cells (Thermo Fisher Scientific) were maintained at 37°C, 5% CO_2_ and 90% humidity in Dublecco’s Modified Eagle Medium (DMEM, Thermo Fisher Scientific) supplemented with 9% fetal bovine serum (FBS) and Penicillin-Streptomycin (both Thermo Fisher Scientific). Tethering experiments were performed in HeLa Tet-Off cells (Clontech) cultured in the same conditions.

### siRNA-mediated knockdowns

The cells were seeded in 6-well plates at a density of 2×10^5^ cells per well and reverse transfected using 2.5 µl Lipofectamine RNAiMAX and 60 pmol of the respective siRNA(s) according to the manufacturer’s instructions. In preparation for mass spectrometry, the cells were reverse transfected in 10 cm dishes using 10 µl Lipofectamine RNAiMAX and 300 pmol siRNA. siRNAs were targeted against Luciferase (5′-CGTACGCGGAATACTTCGA-3′), EIF4A3 (5′-AGACATGACTAAAGTGGAA-3′), RBM8A (5′-TTCGCAGAATATGGGGAAA-3′), CASC3 (5′-CTGATGACATCAAACCTCGAAGAAT-3′, 5′-CGTCATGAACTTTGGTAATCCCAGT-3′), UPF1 (5′-GATGCAGTTCCGCTCCATT-3′), XRN1 (5′-AGATGAACTTACCGTAGAA-3′), SMG6 (5′-GGGTCACAGTGCTGAAGTA-3′) or SMG7 (5′-CGATTTGGAATACGCTTTA-3′).

### Generation of knockout cells using CRISPR-Cas9

The knockouts were performed using the Alt-R CRISPR-Cas9 system (IDT) and reverse transfection of a Cas9:guideRNA ribonucleoprotein complex using Lipofactamine RNAiMAX (Thermo Fisher Scientific) according to the manufacturer’s protocol. The crRNA sequences to target CASC3 were /AlTR1/rGrCrGrCrGrCrUrUrCrGrCrArArGrArCrArCrCrGrGrUrUrUrUrArGrArGrCrUrArUrGrCrU/AlTR2/ (clone H) and /AlTR1/rGrUrUrCrGrGrCrCrUrCrCrGrCrGrCrUrGrUrGrArGrUrUrUrUrArGrArGrCrUrArUrGrCrU/AlTR2/ (clones F and T). Reverse transfection was performed on 1.5×10^5^ cells per crRNA in 12-well dishes. 48 hours after transfection the cells were trypsinized, counted and seeded at a density of a single cell per well in 96-well plates. Cell colonies originating from a single clone were then validated by Sanger sequencing of the targeted genomic DNA locus and western blotting.

### Plasmid transfection

All used plasmids are listed in Supplementary Table S1. To express FLAG-tagged protein constructs and the reporter mRNAs detected by northern blotting, the cells were stably transfected using the Flp-In T-REx system and the tetracycline inducible pcDNA5/FRT/TO vector (Thermo Fisher Scientific). The constructs TPI-WT, TPI-PTC, β-globin WT and β-globin PTC are available on Addgene (IDs 108375-108378). 2.5×10^5^ cells were seeded 24 h before transfection in 6-wells. Per well, 1 µg of reporter construct was transfected together with 1 µg of the Flp recombinase expressing plasmid pOG44 using the calcium phosphate method. 48 h after transfection, the cells were transferred into 10 cm dishes and selected with 100 µg/ml hygromycin. After 10 days, the colonies were pooled. Expression of the reporter mRNA was induced with 1 µg/ml doxycycline for 24 h.

Constructs that express V5-tagged and MS2V5-tagged proteins were stably integrated into the cells using the PiggyBac (PB) Transposon system and the cumate-inducible PB-CuO-MCS-IRES-GFP-EF1-CymR-Puro vector (System Biosciences). 2.5×10^5^ cells were seeded 24 h before transfection in 6-wells. 2.5 µg of the PB Transposon vector and 0.8 µg of PB Transposase were transfected per well using the calcium phosphate method. After 48 h, the cells were pooled in 10 cm dishes and positive clones selected with 2 µg/ml puromycin for a week. Expression of proteins was induced using 30 µg/ml cumate for 72 h.

The tethering construct pSBtet-Hyg-TPI-4MS2-SMG5-4H was stably integrated into HeLa Tet-Off cells using the Sleeping Beauty (SB) transposon system (35, 36). pSBtet-Hyg was a gift from Eric Kowarz (Addgene plasmid #60508; http://n2t.net/addgene:60508; RRID:Addgene_60508). pCMV(CAT)T7-SB100 was a gift from Zsuzsanna Izsvak (Addgene plasmid #34879; http://n2t.net/addgene:34879; RRID:Addgene_34879). 2.5×10^5^ cells were seeded 24 h before transfection in 6-wells. Per well, 1 µg of the reporter construct was transfected together with 1.5 µg of the SB Transposase using the calcium phosphate method. 48 h after transfection, the cells were transferred into 10 cm dishes and selected with 100 µg/ml hygromycin. After 10 days, the colonies were pooled. In absence of tetracycline the reporter was constitutively expressed.

### RNA-Sequencing and computational analyses

RNA-Seq analysis was carried out with 293 wild type (WT) cells transfected with Luciferase siRNA and the CASC3 KO clones H and T transfected with either Luciferase or CASC3 siRNAs. Three biological replicates were analyzed for each sample. RNA was isolated with the kit NucleoSpin RNA Plus (Macherey-Nagel). The Lexogen SIRV Set1 Spike-In Control Mix (SKU: 025.03) that provides a set of external RNA controls was added to the total RNA to enable performance assessment. Mix E0 was added to replicate 1, mix E1 was added to replicate 2 and mix E2 to replicate 3. The Spike-Ins were not used for analysis. The library preparation was performed with the TrueSeq Stranded Total RNA kit (Illumina). First steps of the library preparation involve the removal of ribosomal RNA using biotinylated target-specific oligos combined with Ribo-Zero Gold rRNA removal beads from 1 µg total RNA input. The Ribo-Zero Gold Human/Mouse/Rat kit depletes samples of cytoplasmic and mitochondrial rRNA. Following purification, the RNA is fragmented and cleaved. RNA fragments are copied into first strand cDNA using reverse transcriptase and random primers, followed by second strand cDNA synthesis using DNA Polymerase I and RNase H. These cDNA fragments then have the addition of a single’A’ base and subsequent ligation of the adapter. The products are purified and enriched with PCR to create the final cDNA library. After library validation and quantification (Agilent tape station), equimolar amounts of library were pooled. The pool was quantified by using the Peqlab KAPA Library Quantification Kit and the Applied Biosystems 7900HT Sequence Detection System and sequenced on an Illumina NovaSeq6000 sequencing instrument and a PE100 protocol.

Read processing and alignment was performed as described previously (37). In short, adaptor sequences and low quality bases were removed with Flexbar 3.0 (38). Short reads from the rRNA locus were subtracted by mapping against the 45S precursor (Homo sapiens, NR_046235.1) using Bowtie2 (39). The remaining reads were aligned against the human genome (version 38, EnsEMBL 90 transcript annotations) using the STAR read aligner (version 2.5.3a) (40).

To compute gene differential expression analysis, reads covering exons were counted with FeatureCounts (version 1.5.1) (41) using the ‘—primary’ and ‘—ignoreDup’ parameters. Differential gene expression analysis was performed with DESeq2 (42, 43) and IWH R packages. Significance thresholds were |log2FoldChange|> 1 and adjusted p-value (padj) < 0.05. Genes were designated as small RNA (sRNA) host gene, if they contained other Ensembl-annotated genes of biotypes snoRNA or miRNA within their genomic coordinates (44).

Differential splicing was detected with LeafCutter (version 0.2.7) (45) with the parameters min_samples_per_intron = 2 and min_samples_per_group = 2. Significance thresholds were |deltapsi| > 0.1 and adjusted p-value (p.adjust) < 0.05.

Transcript abundance estimates were computed with Salmon (version 0.13.1) (46) using the the - validateMappings --gcBias parameters. Differential transcript usage was computed with IsoformSwitchAnalyzeR (version 1.7.1) and the DEXSeq method (47–52). Significance thresholds were |dIF| > 0.1 and adjusted p-value (isoform_switch_q_value) < 0.05. For the Boxplot and Kolmogorov-Smirnoff test, the data were filtered only for the adjusted p-value. PTC status of transcript isoforms with annotated open reading frame was determined by IsoformSwitchAnalyzeR using the 50 nt rule of NMD (49,53–55). Isoforms with no annotated open reading frame in Ensembl were designated “NA” in the PTC analysis.

The UPF1 and SMG6/7 (56) and RNPS1 (57) knockdown datasets were processed and analyzed with the same programs, program versions, and scripts as the CASC3 dataset. All packages used are listed in the respective analysis table (Supplementary Tables S4-6). Overlaps of data sets were represented via nVenn (58), eulerr (59) and Upset plots (60). Heatmaps were generated using ComplexHeatmap (61). Barcode plots were produced with barcodeplot function from the limma package version 3.38.3 (50) using transcript isoform dIF as ranking statistic. To test the significance of this enrichment, we used the function cameraPR, from the same package, with the use.rank parameter set to TRUE (62).

### SILAC, co-immunoprecipitation and mass spectrometry

293 WT and 293 CASC3 KO clone H cells expressing either FLAG or FLAG-EIF4A3 were labeled by maintaining them for 5 passages in DMEM for SILAC medium (Thermo Fisher Scientific) supplemented with FBS (Silantes), Penicillin-Streptomycin (Thermo Fisher Scientific) and the respective amino acids at a final concentration of 0.798 mmol/L (Lysine) and 0.398 (Arginine). Unlabeled proline was added to prevent enzymatic Arginine-to-Proline conversion. The conditions were “light” (unlabeled Lysine/Arginine), “medium” (Lysine 4/Arginine 6) and “heavy” (Lysine 8/Arginine 10). A label switch was performed between the three replicates according to the experimental setup listed in Supplementary Table S2. 24 h before expression of the FLAG-tagged construct, the CASC3 KO clone H cells were treated with siRNA against CASC3. The expression of FLAG or FLAG-EIF4A3 was induced for 72 h with 1 µg/ml doxycycline. The cells were lysed in buffer E with RNAse (20 mM HEPES-KOH (pH 7.9), 100 mM KCl, 10% glycerol, 1 mM DTT, Protease Inhibitor, 1 µg/ml RNAse A) and sonicated using the Bandelin Sonopuls mini20 with 15 pulses (2.5 mm tip in 600 µl volume, 1s, 50% amplitude). 600 µl of a 1.6 mg/ml total protein lysate were incubated with 30 µl Anti-FLAG M2 magnetic beads (Sigma) at 4°C while rotating for 2 h. The beads were washed three times for 5 min with EJC-buffer (20 mM HEPES-KOH (pH 7.9), 137 mM NaCl, 2 mM MgCl_2_, 0.2% Triton X-100, 0.1% NP-40) and eluted in 43 µl of a 200 mg/ml dilution of FLAG peptides (Sigma) in 1x TBS. The samples were merged according to Supplementary Table S2. 1 volume of 10% SDS was added and the samples were reduced with DTT and alkylated with CAA (final concentrations 5 mM and 40 mM, respectively). Tryptic protein digestion was performed using a modified version of the single pot solid phase-enhanced sample preparation (SP3) (63). In brief, reduced and alkylated proteins were supplemented with paramagnetic Sera-Mag speed beads (Thermo Fisher Scientific) and mixed in a 1:1-ratio with 100% acetonitrile (ACN). After 8 min incubation protein-beads-complexes were captured using an in-house build magnetic rack and two times washed with 70% EtOH. Afterwards, samples were washed once with 100% ACN, air-dried and reconstituted in 5 µl 50 mM Triethylamonium bicarbonate supplemented with 0.5 µg trypsin and 0.5 µg LysC and incubated overnight at 37°C. On the next day, the beads were resuspended and mixed with 200 µl ACN, incubated for 8 min and again placed on the magnetic rack. Tryptic peptides were washed once with 100% ACN, airdried, dissolved in 4% DMSO and transferred into 96-well PCR tubes. After acidification with 1 µl of 10% formic acid, the samples were ready for LC-MS/MS analysis.

Proteomics analysis was performed by data-dependent acquisition using an Easy nLC1200 ultra high-performance liquid chromatography (UHPLC) system coupled via nanoelectrospray ionization to a Q Exactive Plus instrument (all Thermo Scientific). Tryptic peptides were separated based on their hydrophobicity using a chromatographic gradient of 60 min with a binary system of buffer A (0.1% formic acid) and buffer B (80% ACN, 0.1% formic acid). In-house made analytical columns (length: 50 cm, inner diameter: 75 µm) filled with 1.9 µm C18-AQ Reprosil Pur beads (Dr. Maisch) were used for separation. Buffer B was linearly increased from 3% to 27% over 41 min followed by a steeper increase to 50% within 8 min. Finally, buffer B was increased to 95% within 1 min and stayed at 95% for 10 min in order to wash the analytical column. Full MS spectra (300 – 1,750 m/z) were acquired with a resolution of 70,000, a maximum injection time of 20 ms, and an AGC target of 3e6. The top 10 most abundant peptide ions of each full MS spectrum were selected for HCD fragmentation (NCE: 27) with an isolation width of 1.8 m/z and a dynamic exclusion of 10 seconds. MS/MS spectra were measured with a resolution of 35,000, a maximum injection time of 110 ms and an AGC target of 5e5.

MS RAW files were analysed using the standard settings of the MaxQuant suite (version 1.5.3.8) with the before mentioned SILAC labels (64). Peptides were identified by matching against the human UniProt database using the Andromeda scoring algorithm (65). Carbamidomethylation of cysteine was set as a fixed modification, methionine oxidation and N-terminal acetylation as variable modification. Trypsin/P was selected as the digestion protein. A false discovery Rate (FDR) < 0.01 was used for identification of peptide-spectrum matches and protein quantification. Data processing and statistical analysis was done in the Perseus software (version 1.5.5.3) (66). Significantly changed proteins were identified by One-sample t-testing (H0 = 0, fudge factor S0 = 0.1). The results are listed in Supplementary Table S2. Visualization was performed with the Instant Clue software (version 0.5.3) and the R package ggplot2 (version 3.1.0) (67, 68).

Co-immunoprecipitation experiments followed by western blotting were performed as described above except that a 15 min incubation step in SDS buffer (600 mM Tris pH 6.8, 100 mM DTT, 10% Glycerol, 2% SDS, 0.002% Bromophenolblue) was used for elution from the beads.

### Semi-quantitative and quantitative reverse transcriptase (RT)-PCR

RNA was extracted using peqGOLD TriFast reagent (VWR) according to the manufacturer’s instructions. Reverse transcription was performed with GoScript Reverse Transcriptase (Promega) using 2 µg total RNA and oligo dT primers. Semi-quantitative PCR was carried out with MyTaq Red Mix (Bioline). Quantitative real time PCR was performed with 16 ng of cDNA per reaction with GoTaq qPCR Master Mix (Promega) and the CFX96 Touch Real-Time PCR Detection System (Biorad). The average cT values were calculated from three technical replicates. The mean fold changes from three biological replicates were calculated according to the ΔΔCt method (69). When measuring isoform switches, the fold change of the PTC-containing transcript was normalized to the canonical transcript. When measuring differential expression, the fold change was normalized to GAPDH. For each primer pair amplification efficiencies were measured by a 2-fold dilution curve and ranged between 87 and 100.1%. The primer sequences are listed in Supplementary Table S3.

### Western blotting

Protein extraction was performed with peqGOLD TriFast reagent (VWR), and proteins were separated by SDS-PAGE gel electrophoresis and transferred to a PVDF membrane (GE Healthcare Life Sciences). The following antibodies were used: anti-CASC3 amino acid residues 653-703 (Bethyl Laboratories, #A302-472A-M), anti-CASC3 amino acid residues 367-470 (Atlas Antibodies, #HPA024592), anti-EIF4A3 (Genscript), anti-FLAG (Cell Signaling Technology, #14793), anti-RBM8A (Atlas Antibodies, #HPA018403), anti-SMG6 (Abcam, #ab87539), anti-SMG7 (Elabscience, #E-AB-32926), anti-Tubulin (Sigma-Aldrich, #T6074), anti-V5 (QED Bioscience, #18870), anti-XRN1 (Bethyl Laboratories, #A300-443A), anti-rabbit-HRP (Jackson ImmunoResearch, #111-035-006), anti-mouse-HRP (Jackson ImmunoResearch, #115-035-003). Detection was performed with Western Lightning Plus-ECL (PerkinElmer) or Amersham ECL prime (Ge Healthcare Life Sciences) and the chemiluminescence imager Fusion FX6 EDGE (Vilber-Lourmat).

### Northern blotting

The cells were harvested in peqGOLD TriFast reagent (VWR) and total RNA extraction was performed as recommended by the manufacturer’s protocol. 2.5 μg of total RNA were resolved on a 1% agarose/0.4 M formaldehyde gel using the tricine/triethanolamine buffer system (70) followed by a transfer on a nylon membrane (Roth) in 10x SSC. The blots were incubated overnight at 65°C in Church buffer containing [α-32P]-GTP body-labeled RNA probes for detection of the reporter mRNA. Endogenous 7SL RNA was detected by a 5′-32P-labeled oligonucleotide (5′-TGCTCCGTTTCCGACCTGGGCCGGTTCACCCCTCCTT-3′). The blots were visualized and quantified using a Typhoon FLA 7000 (GE Healthcare) and ImageQuant TL 1D software.

## Results

### CASC3 is dispensable for the correct splicing of many EJC-dependent transcripts

Previously, we and others have shown that the depletion of EJC core components in human cells leads to pervasive re-splicing of cryptic splice sites, resulting in aberrant splice variants lacking exonic sequences (37, 71). Mechanistically, the EJC prevents the use of cryptic splice sites either by interaction with the ASAP/PSAP component RNPS1 or by sterically masking splice sites (schematically depicted in Supplementary Figure S1A) (37). In keeping with previous observations from HeLa cells, the knockdowns of the EJC core components EIF4A3 or RBM8A in HEK293 cells resulted in exon-skipping of the mRNAs for RER1, OCIAD1 and MRPL3 (Figure 1A-C). Surprisingly, when we performed a knockdown of the EJC core factor CASC3 this did not result in mis-splicing of these three selected transcripts (Figure 1A-C). This observation stands in contrast to a previous study that has reported transcriptome-wide alternative splicing upon CASC3 depletion, including MRPL3 (71, 72). Interestingly, the knockdown of CASC3 increased the abundance of an alternative transcript isoform of the SR protein SRSF2 (Figure 1D), which was previously shown to be regulated by NMD. Although this altered abundance of the NMD-targeted SRSF2 transcript isoform could be due to alternative splicing, we consider it very likely that reduced NMD is responsible for this effect.

**Figure 1.**
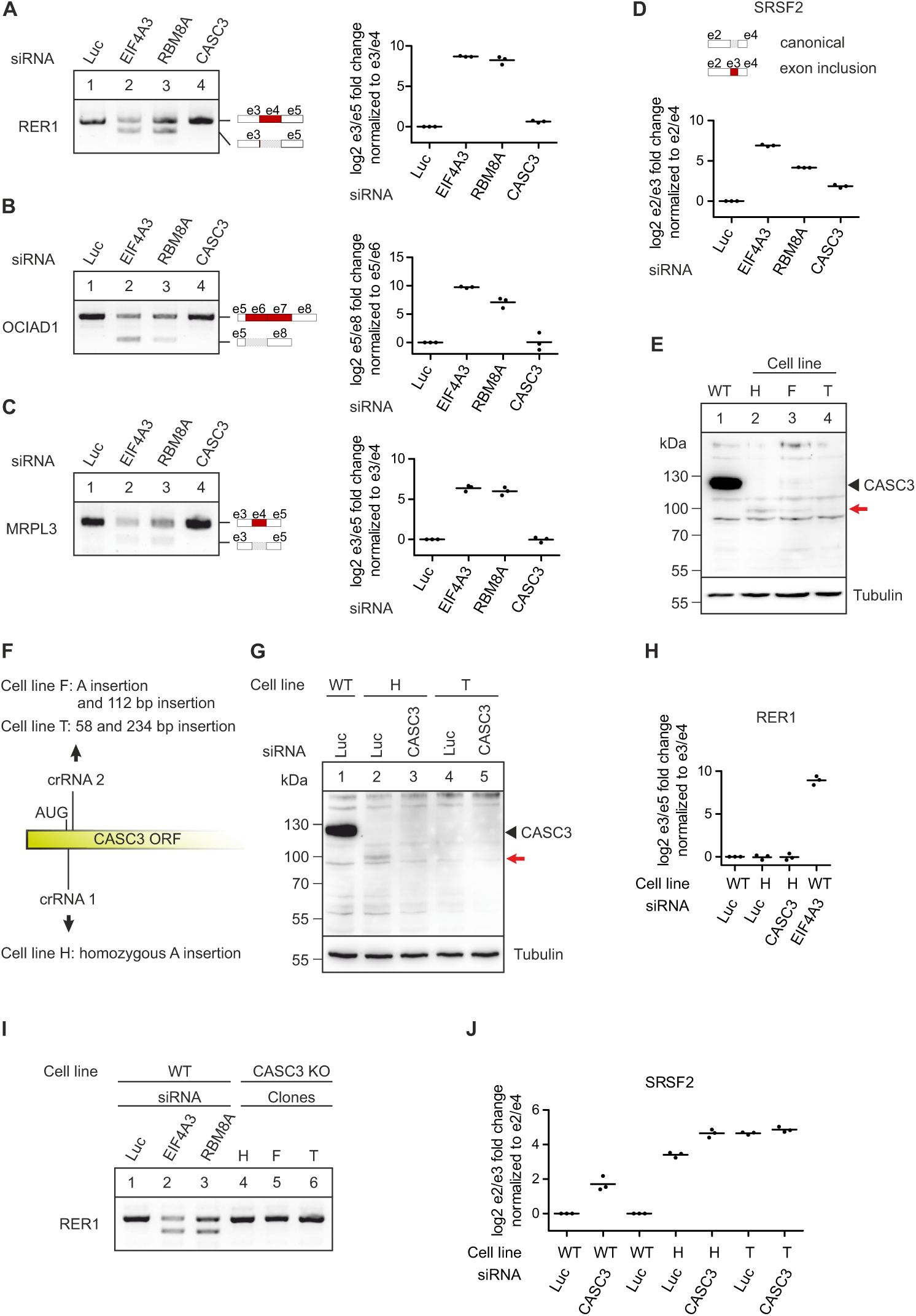
CASC3 is not involved in the splicing regulation of known EJC-dependent targets. **A-C**: RT-PCR- and quantitative RT-PCR-based detection (qPCR) of transcript isoforms of the genes RER1 (A), OCIAD1 (B), and MRPL3 (C) after siRNA-mediated knockdown of the indicated EJC components or Luciferase (Luc) as a negative control. Skipped exons are depicted schematically (e: exon). Data points and means from the qPCRs are plotted (n=3). **D**: Relative quantification of the SRSF2 transcript isoforms by qPCR following knockdown of the indicated EJC components or Luciferase (Luc) as a negative control. The transcript variants at the position of the included exon are depicted schematically. Data points and means are plotted (n=3). **E**: Total protein lysates from wild-type cells (WT) and CASC3 knockout (KO) cell lines H, F and T were separated by SDS-PAGE and CASC3 was detected by western blotting. The red arrow indicates an additional band detected in the cell lines H and F and which is not visible in the WT condition or in the cell line T. **F**: Schematic depiction of the insertions resulting in a CASC3 KO or constitutive knockdown in the indicated clones. **G**: Total protein lysates from WT and CASC3 KO cell lines H and T were separated by SDS-PAGE and CASC3 was detected by western blotting. In lanes 3 and 5 the cells have additionally been treated with siRNAs targeting CASC3. **H**: RT-PCR of transcript isoforms of the gene RER1 after siRNA-mediated knockdown of the indicated EJC components or Luciferase (Luc) as a negative controle, compared to CASC3 KO cell lines H, F and T. **I**: Relative quantification of the RER1 transcript isoforms by qPCR in WT cells treated with Luc siRNA as a negative control, CASC3 KO cell line H treated with Luc siRNA, CASC3 KO cell line H treated with CASC3 siRNAs and WT cells treated with EIF4A3 siRNA. Data points and means are plotted (n=3). **J:** Relative quantification of the SRSF2 transcript isoforms by qPCR in WT cells treated with Luc siRNA as a negative control, WT cells treated with CASC3 siRNA, CASC3 KO cell line H treated with Luc siRNA, CASC3 KO cell line H treated with CASC3 siRNAs and CASC3 KO cell line T treated with Luc siRNA as well as CASC3 KO cell line T treated with CASC3 siRNAs. Data points and means are plotted (n=3).

Although the knockdown efficiency of CASC3 was substantial (Supplementary Figure S1B), we wished to exclude that residual amounts of CASC3 prevented a reliable assessment of the protein’s function. Since CASC3 was found to be non-essential in multiple genome-wide screens of human immortalized cell lines, we reasoned that knockout (KO) of CASC3 should be feasible (73, 74). Accordingly, we obtained three cell lines by CRISPR-Cas9-mediated gene editing, designated H, F, and T lacking the CASC3-specific 130 kDa band on a western blot (Figure 1E). In all cell lines we detected genomic insertions of different length and sequence at the beginning of the coding region of CASC3, which resulted in frame shifts of the downstream coding region or, in the case of cell line T, contained in-frame termination codons (Figure 1F, Supplementary Figure S1C and D). For the cell lines H and F, we observed an additional band of 100 kDa on western blots with antibodies recognizing the C-terminal or central region of CASC3 (Figure 1E, red arrow, Supplementary Figure S1C and E). This cross-reactive protein interacted with FLAG-tagged EIF4A3 and disappeared upon treatment with siRNAs against CASC3 (Figure 1G, Supplementary Figure S1E and F). This suggests that the cell lines H and F produce an N-terminally truncated form of CASC3, presumably by initiating protein translation at a non-canonical initiation codon. The production of aberrant protein forms by alternative translation initiation has been recently described in a systematic analysis to commonly occur in KO cell generated by CRISPR-Cas9 genome editing (75, 76). The cell line T (without further treatment) and cell line H in combination with CASC3 siRNA treatment completely lack detectable CASC3 protein. This set of cell lines therefore enables a hitherto unfeasible analysis of CASC3’s cellular function.

In agreement with the data obtained from the knockdown of CASC3, transcripts containing EJC-dependent splice sites were correctly spliced in CASC3 KO cells (Figure 1H and I, Supplementary Figure S1G and H). Remarkably, the increased abundance of the NMD-regulated SRSF2 transcript isoform was much more prominent in the CASC3 KO than in the CASC3 knockdown (Figure 1J).

### CASC3 regulates NMD-sensitive isoforms

So far, our results have indicated that CASC3 shapes the transcriptome differently compared to EIF4A3, MAGOH and RBM8A. To investigate the global effects of CASC3 depletion on the transcriptome, we performed RNA-sequencing (RNA-seq) of the cell lines H and T either treated with CASC3 or control siRNAs (Figure 2A) and identified differentially expressed genes (Supplementary Figure S2A and B, Supplementary Table S4). To exclude clone-specific and siRNA treatment-related effects, we compared the identified targets between the four conditions. Overall, the high number and the substantial overlap of upregulated genes suggests that CASC3 KO mainly results in the accumulation of certain transcripts (Figure 2B and C, Supplementary Figure S2C). Interestingly, several upregulated genes belong to the class of small RNA (e.g. snoRNA) host genes, which are frequently NMD targets (Figure 2C and D) (31, 33). We validated the upregulation of the snoRNA host gene ZFAS1 by qPCR, which was even more pronounced in CASC3 KO than UPF1 knockdown cells (Supplementary Figure S2D and E). Across the top 100 significantly upregulated genes in CASC3 KO, 15 small RNA host genes were identified (Figure 2D). Comparing the differentially expressed genes to recent transcriptome-wide NMD screens, many of the top 100 significantly upregulated genes were also differentially expressed in UPF1 and SMG6/7 KD (Figure 2D, 23% and 59%, respectively) (31, 56). However, none of the identified targets were present in an RNPS1 knockdown (Figure 2D) (37, 57). Collectively, our differential gene expression analysis strengthens the proposed link between CASC3 and the NMD-machinery.

**Figure 2.**
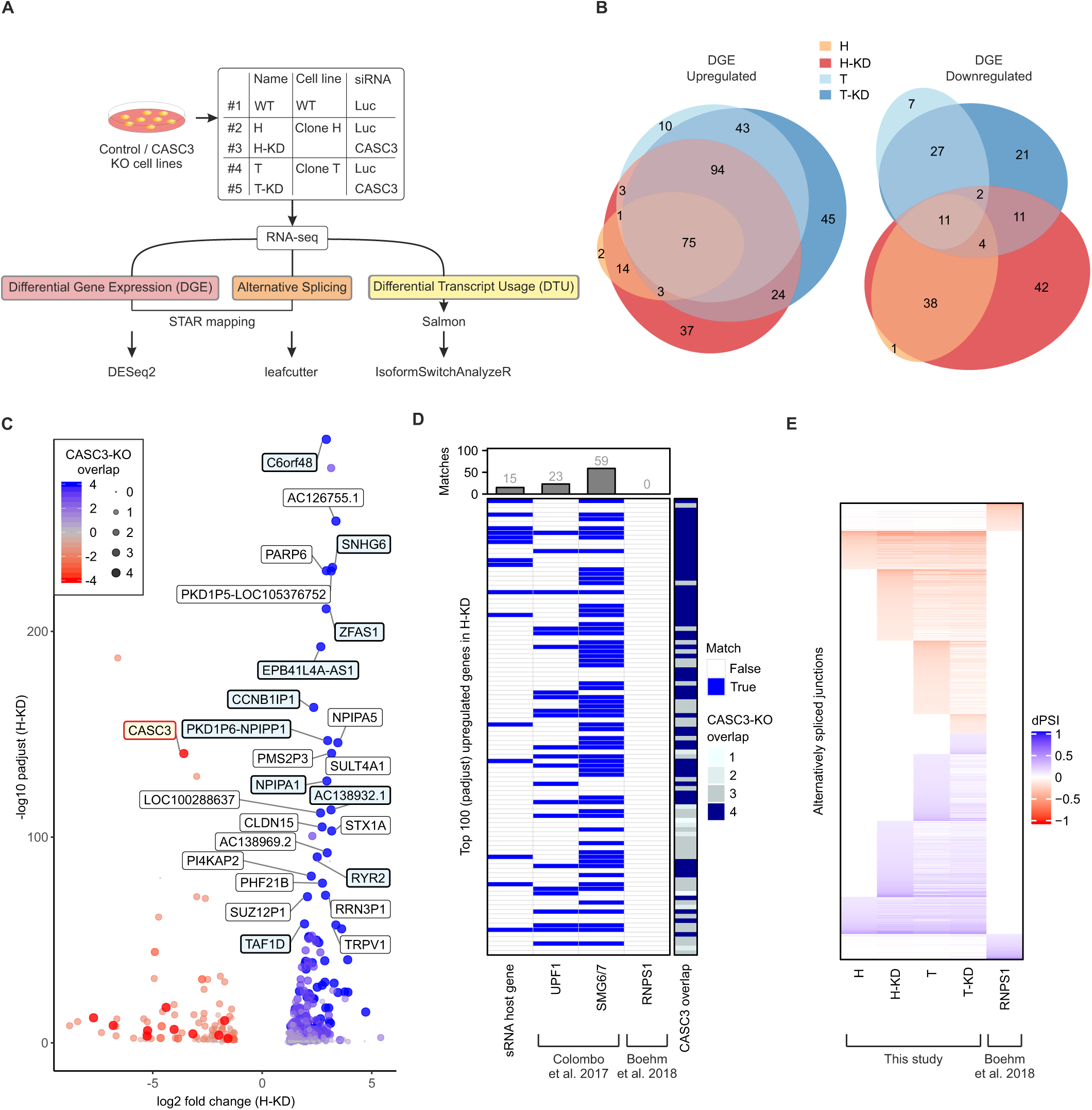
Transcriptome-wide effects of CASC3 depletion. **A**: Workflow for RNA-sequencing analysis. **B**: Overlap of up- and downregulated genes in the CASC3 KO cell lines H and T, +/- CASC3 siRNAs. DGE: Differential gene expression. Due to the visualization as an Euler plot, some intersections cannot be plotted. All intersections are shown in Supplementary Figure S2C. **C**: Volcano plot of differential gene expression analysis of the condition H-KD using overlap from Figure 2B as color and point size definition. Gene symbols are indicated for the top 25 upregulated genes detected in all four conditions and for CASC3 (colored in light red). Labels of small RNA host genes are colored in light blue. Log2 fold change is plotted against –log10 padjust (adjusted p-value). **D**: Matching of the top 100 upregulated genes sorted by padjust (adjusted p-value) in condition H-KD with small RNA (sRNA) host genes and comparison to knockdowns of UPF1, SMG6/7 and RNPS1. **E**: Heatmap of all identified alternatively spliced junctions in the respective condition, measured in delta percent spliced in (dPSI).

Next, we analyzed alternative splicing changes in CASC3 KO cells (Figure 2E). Since our earlier assays showed that CASC3 was not involved in the EJC-regulated splicing of many targets (Figure 1), we were surprised to detect many altered splicing events in CASC3 depleted cells (Supplementary Figure S2F, Supplementary Table S5). It is remarkable that hardly any alternative splicing events were shared between RNPS1 knockdown and CASC3 KO cells (Figure 2E, Supplementary Figure S2F). Either CASC3 regulates an RNPS1-independent set of alternative splice sites or the splicing changes are due to impaired NMD, which fails to remove NMD-sensitive isoforms. To test these possibilities, we investigated the functional consequence of CASC3-dependent alternative splicing on the transcript isoform level (Figure 3A, Supplementary Figure S3A and B, and Supplementary Table S6). Strikingly, in all CASC3 KO conditions many upregulated mRNA isoforms contained a premature termination codon (PTC), rendering the transcripts susceptible to NMD (Figure 3A). On the other hand, downregulated isoforms rarely contained a PTC. Among the identified isoform switches was the target SRSF2, which we confirmed earlier to be CASC3-dependent (Figure 1D and J). While overall SRSF2 gene expression varied only slightly between wild-type and CASC3 KO cells, the isoform usage changed dramatically towards the accumulation of NMD-sensitive transcripts in the CASC3 KO conditions (Figure 3B). A similar accumulation of NMD-sensitive transcripts was also observed for other transcripts (Supplementary Figure S3C-E).

**Figure 3.**
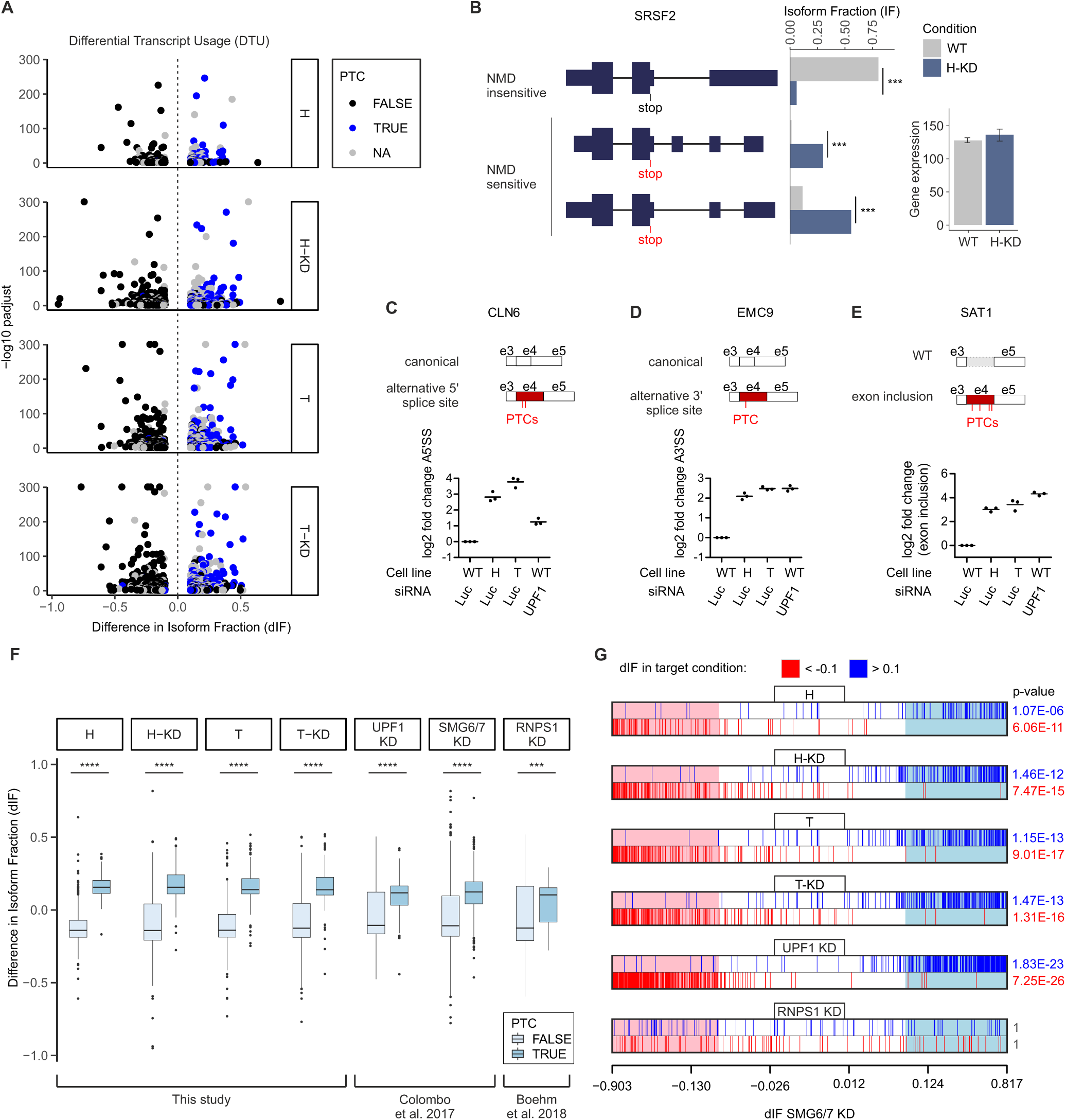
Knockout of CASC3 leads to a global upregulation of NMD-sensitive transcript isoforms. **A**: Results from IsoformSwitch analysis plotted as volcano plots. Transcript isoforms identified as NMD-sensitive are shown as blue dots. Isoforms with no annotated open reading frame are designated as “NA”. Difference in Isoform Fraction (dIF) is plotted against –log10 padjust (adjusted p-value). **B**: Quantification of transcript isoforms from SRSF2 by IsoformSwitchAnalyzeR. **C-E**: Relative quantification of the schematically depicted transcript isoforms of the genes CLN6 (C), EMC9 (D), and SAT1 (E) by qPCR in WT cells, CASC3 KO cell lines H and T and WT cells treated with siRNA targeting UPF1. PTC: premature termination codon. Individual data points and means are plotted (n=3). **F**: Boxplot of PTC-containing vs. non-PTC-containing transcript isoforms after IsoformSwitch analysis for all CASC3 KO conditions compared to UPF1, SMG6/7 and RNPS1. A Kolmogorov-Smirnoff test was applied (p-value < 0.001 ***, p-value < 10-16 ****). **G**: Barcode plots showing the enrichment of transcript isoforms that undergo isoform switching (padj < 0.05) of a target condition compared to the SMG6/7 KD dataset. On the x-axis the dIF of transcripts that undergo isoform switching in SMG6/7 are ranked according to their dIF. The regions in the barcode plot with dIF > 0.1 of SMG6/7 KD are shaded light red if dIF < −0.1 and light blue if dIF > 0.1. Similarly, individual transcript isoforms with a |dIF| > 0.1 in the target conditions are marked with red lines if |dIF| < −0.1 and blue lines if dIF > 0.1. To test whether the up- and downregulated sets of transcripts in the target conditions are highly ranked in terms of dIF relative to transcripts that are not in the respective set, the camera test from the limma R package was performed and significant p-values (< 0.05) are shown in blue (up) or red (down).

We next validated a set of transcript isoform switches by qPCR (Figure 3C-E, Supplementary Figure S3F and G). In the transcript isoforms stabilized by the CASC3 KO and by a UPF1 KD, the inclusion of intronic regions resulted in the inclusion of PTCs. The shift of isoform usage from NMD-insensitive to PTC-containing transcripts was also observed transcriptome-wide in CASC3 KO cells and was comparable to NMD-compromised SMG6/7 or UPF1 depleted cells (Figure 3F). When compared to transcript isoform changes in a SMG6/7 knockdown, the events in the CASC3 KO are enriched in a similar fashion (Figure 3G). While the enrichment is stronger in a comparison of UPF1 and SMG6/7, the isoform changes occurring in the RNPS1 KD do not correlate with the ranked SMG6/7 events (Figure 3G, Supplementary Figure S3H). These findings indicate that many transcript isoforms upregulated upon depletion of CASC3 represent genuine endogenous NMD targets.

### The EJC core is undisturbed if CASC3 is not present

The CASC3 KO could potentially influence the composition of exon junction complexes and their peripheral interacting proteins, which could underlie the observed effects on the transcriptome. Therefore, we analyzed the FLAG-tagged EIF4A3 interactome in the cell line H treated with CASC3 siRNAs and wild type cells using mass spectrometry. EIF4A3 was successfully enriched, together with other known EJC complex members (Figure 4A and B, Supplementary Figure S4). Co-precipitated CASC3 was reduced to background levels in the knockout condition, further validating the absence of CASC3 in the EJC (Figure 4B and C, log2 fold change = −4.34). We were interested to identify factors that significantly changed between the KO and WT condition. These belong to three distinct groups: significant in both WT/CTL and KO/CTL conditions (blue dots), significant in KO/CTL (orange dots), and significant in WT/CTL (magenta dots). In the first group, no factor was changed between the WT and KO condition more than 1.6 fold (Supplementary Figure S4). In the second group three factors were negatively enriched in the KO/CTL condition or marginally altered between KO and WT conditions (Figure 4B and C, WARS, CMBL and USP15).

**Figure 4.**
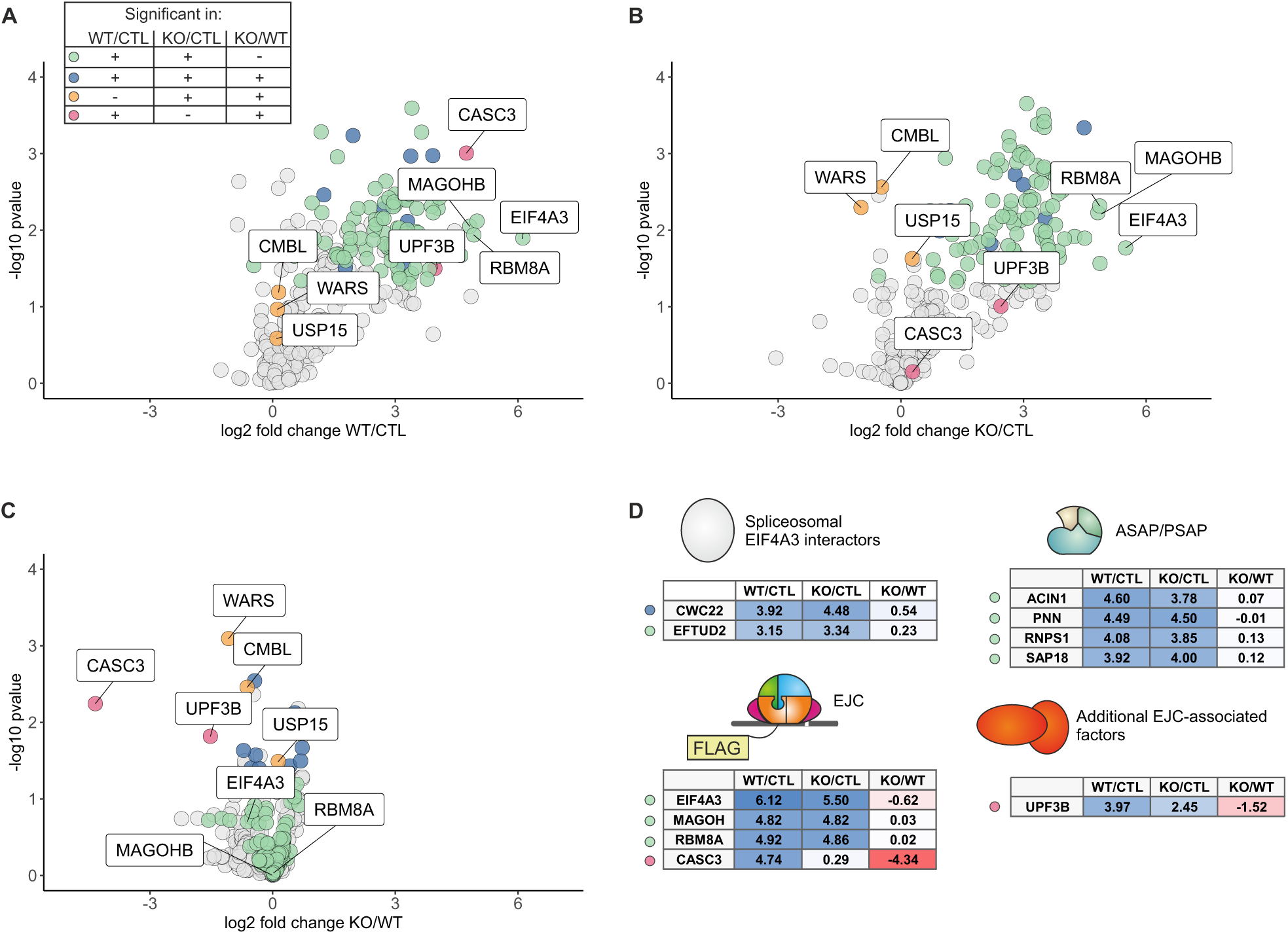
Cells that lack CASC3 have intact EJCs. **A-C**: Volcano plots of mass spectrometry-based analysis of the interaction partners of EIF4A3 in WT cells and in the CASC3 KO cell line H treated with siRNAs targeting CASC3. A: EIF4A3 against FLAG control in WT cells, B: EIF4A3 against FLAG control in KO cells, C: EIF4A3 in KO cells against EIF4A3 in WT cells. The color labeling indicates targets that are significant in the respective comparisons after one-sample t-testing. **D**: Overview of the enrichment of EJC- and EJC-associated proteins.

The only other protein besides CASC3 that was significantly changed between KO/WT and only enriched in the WT/CTL condition was the NMD factor UPF3B (Figure 4C, log2 fold change = −1.52). UPF3B links the EJC to the NMD machinery via direct interactions (77) and was recently found to be enriched in cytoplasmic CASC3-loaded EJCs (22). The reduction of NMD-competent EJCs could contribute to the NMD impairment that we observed upon loss of CASC3. Strikingly, no other EJC core factor or splicing regulatory EJC component (e.g. ASAP/PSAP) was considerably altered in the CASC3-depleted condition (Figure 4D).

### CASC3 stimulates SMG6-dependent endonucleolytic cleavage

To deepen the understanding of how a lack of CASC3 results in reduced NMD efficiency, we stably integrated the well-established globin NMD reporter PTC39 in WT and CASC3 KO cell lines (Figure 5A and B, Supplementary Figure S5A) (78). The analysis of a reporter mRNA enables a read-out of multiple aspects of mRNA degradation: firstly, the total levels of the full-length reporter, secondly the contribution of 5′->3′ exonucleolytic decay by XRN1 (detection of xrFrag due to an XRN1-resistant element) (79, 80); and thirdly the amount of endonucleolytic cleavage by SMG6 (detection of 3′ fragment stabilized by XRN1 knockdown). In both WT and CASC3 KO cell lines the reporter was efficiently degraded, showing that the NMD pathway is still functional in CASC3 depleted cells. However, full-length reporter levels in CASC3 KO were slightly higher when compared to wild-type cells (lane 2 vs. lane 5). Notably, the accumulation of 3′ fragments following XRN1 knockdown was clearly reduced in the CASC3 KO condition (lane 3 vs. lane 6). This difference in endonucleolytic cleavage efficiency was also observed when expressing a minigene reporter of the endogenous CASC3 target TOE1. While there was a substantial upregulation of full-length reporter mRNA in CASC3 KO cells, the amount of the 3′ fragment was strongly reduced, suggesting that SMG6-dependent endonucleolytic cleavage is inefficient in CASC3 KO cells (Figure 5C-E, Supplementary Figure S5B and C). To further address this, a TPI reporter was expressed in combination with knockdowns of XRN1, SMG6 and/or SMG7 (Figure 5F and G, Supplementary Figure S5D). The degree of reporter and 3′ fragment stabilization of the TPI reporter following XRN1 knockdown was comparable to the observations made for the globin and TOE1 reporters (lanes 1-3 vs. lanes 6-8). In both WT and CASC3 KO cells, the SMG6 knockdown resulted in a drastic reduction of 3′ fragments, as expected (lanes 4 and 9). Notably, a knockdown of SMG7 together with XRN1 revealed a major difference between the cell lines. In WT cells the PTC-containing reporter was only minimally stabilized by the SMG7/XRN1 knockdown and 3′ fragments were unaffected. Performing a SMG7/XRN1 knockdown in CASC3 KO cells lead to a more dramatic stabilization of the full-length reporter and a decrease of 3′ fragments compared to the XRN1 knockdown condition (lanes 3 and 5 vs. lanes 8 and 10). Collectively, our results indicate that in CASC3 KO cells SMG6-mediated endonucleolytic cleavage is impaired. This could explain why the CASC3 KO cells are more sensitive to a knockdown of SMG7 when compared to wild type cells.

**Figure 5.**
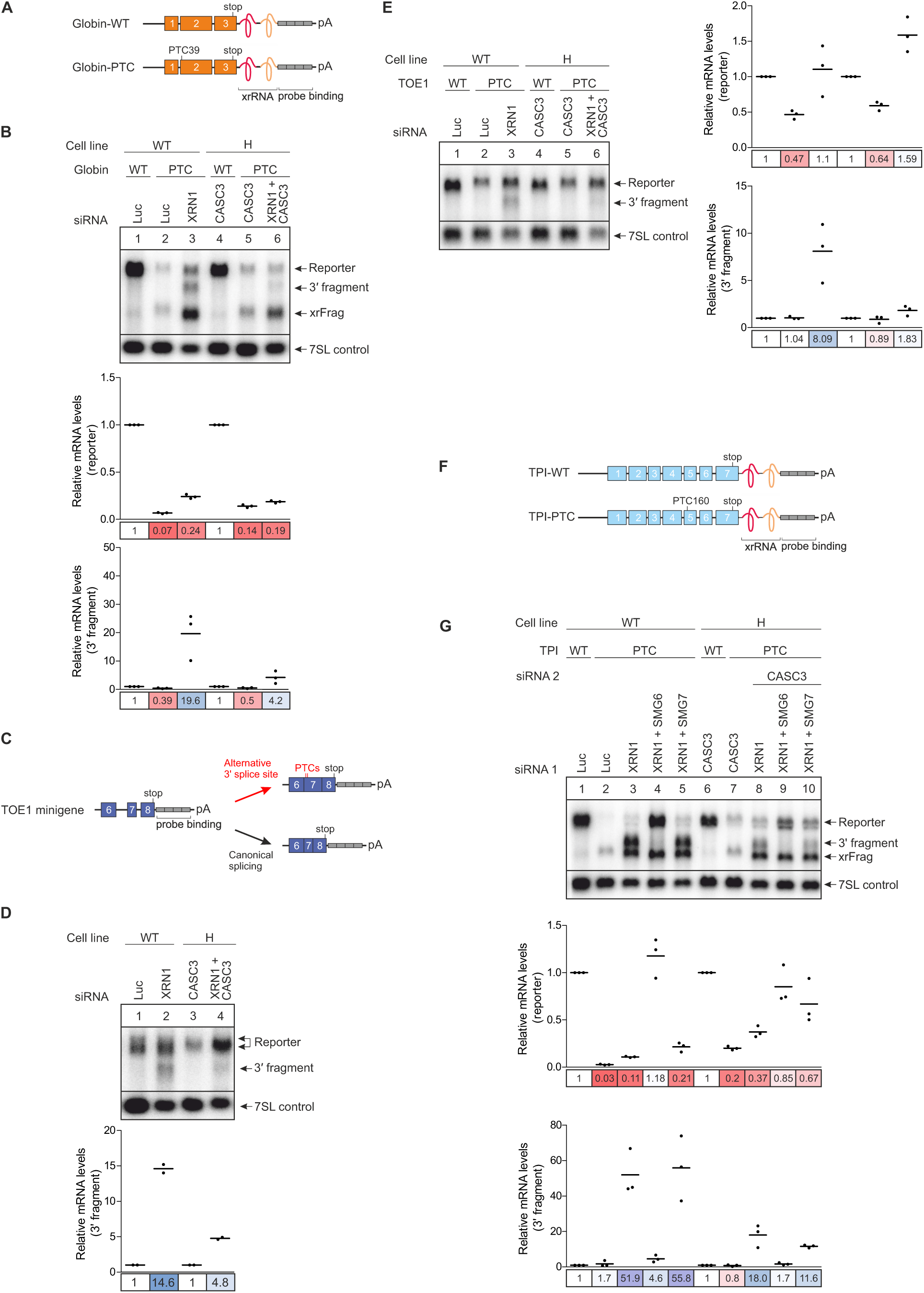
SMG6-mediated endocleavage is impaired when CASC3 is not present. **A**: Schematic depiction of the globin mRNA reporter. The reporter consists of three exons (orange boxes) followed by an XRN1-resistant element (xrRNA) and a probe binding cassette (gray boxes). The PTC reporter contains a premature termination codon (PTC) in the second exon. **B**: Northern blot of RNA extracted from the indicated cell lines that stably express the globin reporter. The xrFrag corresponds to the 3′ part of the reporter that is resistant to degradation by XRN1 due to the xrRNA. The cell lines in lane 3 and 6 were additionally treated with XRN1 siRNA which results in the appearance of a 3′ degradation fragment below the full-length reporter. Reporter and 3′ fragment mRNA levels were normalized to 7SL RNA which is shown as a loading control. For the relative mRNA quantification, in each condition (WT vs. CASC3 KO with KD) the reporter and 3′ fragment levels were normalized to the globin WT reporter (lanes 1 and 4). Individual data points and means are plotted from n=3 experiments. **C**: Schematic depiction of the TOE1 minigene reporter consisting of exons 6-8 (purple boxes) followed by a probe binding cassette (gray boxes). The reporter can be spliced to either contain the canonical stop codon (bottom right) or, by usage of an alternative 3′ splice site, a PTC in exon 7 (top right). **D**: Northern blot of RNA extracted from the indicated cell lines treated with the indicated siRNAs stably expressing the TOE1 minigene reporter. The 3′ fragment levels were first normalized to the 7SL RNA loading control and for every cell line the XRN1 knockdown condition to the condition without XRN1 knockdown (n=2). **E**: Northern blot of RNA extracted from the indicated cell lines treated with the indicated siRNAs stably expressing a prespliced variant of the TOE1 minigene reporter depicted in Figure 5C. The intron between exon 6 and 7 is deleted so that the reporter is constitutively spliced to contain either a normal stop codon (TOE1-WT) or a PTC (TOE1-PTC). The mRNA levels were normalized to 7SL RNA. For the relative mRNA quantification, in each condition (WT vs. CASC3 KO with KD) the reporter and 3′ fragment levels were normalized to the TOE-WT reporter (lanes 1 and 4). **F**: Schematic depiction of the triose phosphate isomerase (TPI) mRNA reporter. The reporter consists of seven exons (blue boxes) followed by an XRN1-resistant element (xrRNA) and a probe binding cassette (gray boxes). The PTC reporter contains a premature termination codon (PTC) in the fifth exon. **G**: Northern blot of RNA extracted from the indicated cell lines treated with the indicated siRNAs stably expressing the either the TPI WT or TPI PTC mRNA reporter. The reporter and 3′ fragment mRNA levels were normalized to the 7SL control. For each cell line, the mRNA levels were then normalized to the respective TPI WT reporter or 3′ fragment levels. Individual data points and means are plotted from n=3 experiments.

To identify, which part of CASC3 promotes NMD, we employed a tethering reporter that was designed to monitor mRNA turnover as well as endonucleolytic cleavage at the termination codon (Figure 6A). Tethering the full-length CASC3 protein to the MS2 stem loops downstream of the stop codon resulted in degradation of the reporter compared to tethering of the negative control GST, as was previously reported for similar tethering reporters (Figure 6B and C, Supplementary Figure S6) (12, 32). This degradation was accompanied by the production of 3′ fragments in XRN1 knockdown conditions, indicating that the mechanism of decay is comparable to the PTC-containing reporter mRNAs (Figure 6B, lane 6). Surprisingly, C-terminally truncated deletion mutants of CASC3 that contain the first 480 or even 137 amino acid residues were able to induce degradation of the tethering reporter to a comparable extent as full-length CASC3 (Figure 6B, lanes 3 and 4, 7 and 8).

**Figure 6.**
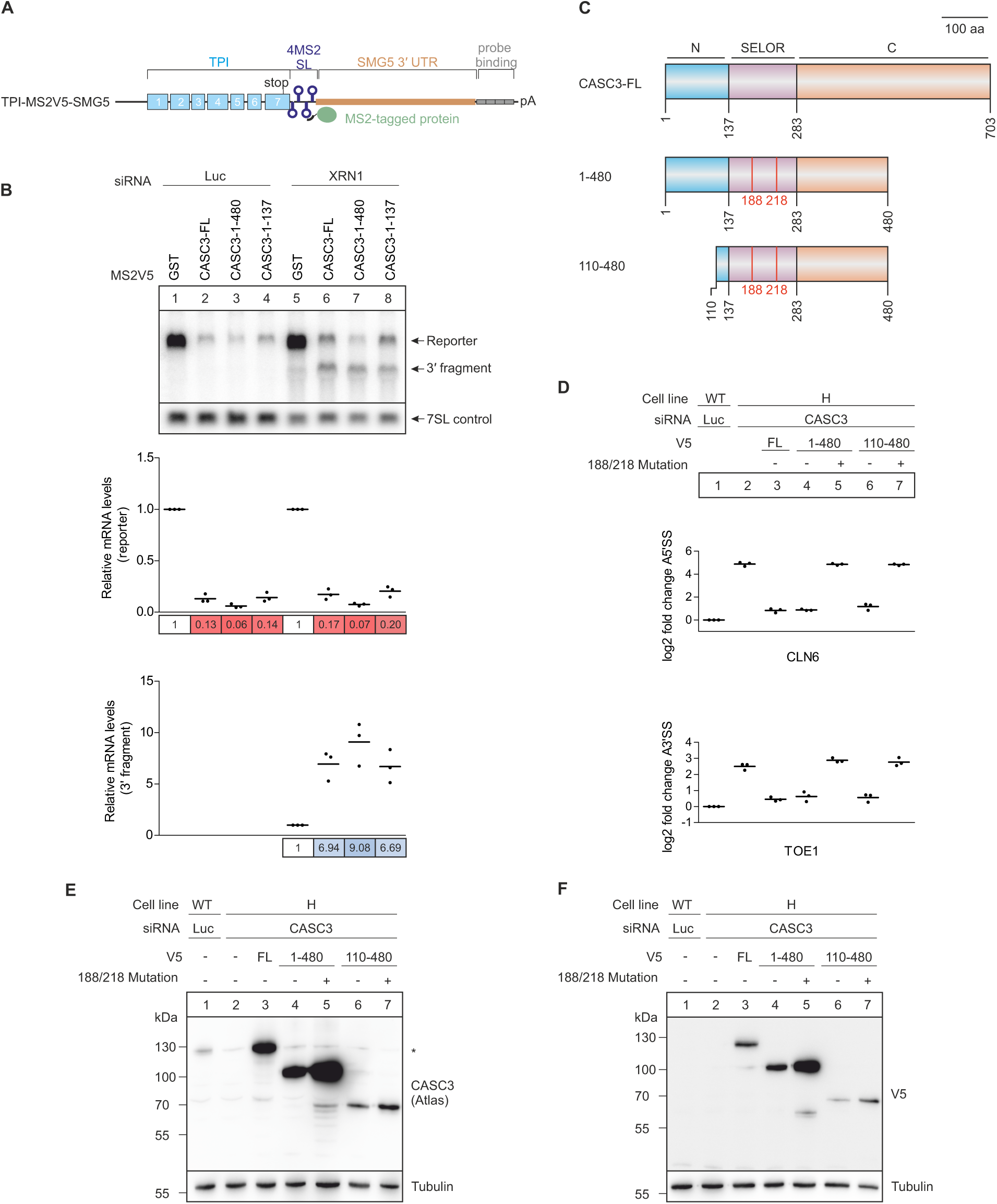
The CASC3 N-terminus promotes but is not necessary to elicit NMD. **A**: Schematic depiction of the TPI-MS2V5-SMG5 tethering reporter. The reporter consists of the TPI ORF (blue boxes) followed by 4 MS2 stem loops (SL). Downstream the SMG5 3′ untranslated region (UTR) is inserted to increase the size of 3′ fragments that result from cleavage at the termination codon. Reporter and 3′ fragment mRNAs can be detected via the probe binding cassette (gray boxes). **B**: Northern blot of a tethering assay performed in HeLa Tet-Off cells. The cells stably express the tethering reporter shown in Figure 6A together with the indicated MS2V5-tagged proteins. When the cells are additionally treated with XRN1 siRNA, a 3′ degradation fragment can be detected below the full-length reporter. The reporter and 3′ fragment mRNA levels are normalized to the 7SL RNA. For the calculation of the relative mRNA levels in each condition (Luc vs. XRN1) the levels were normalized to the MS2V5-GST control (lanes 1 and 5). **C**: Schematic depiction of CASC3 rescue protein constructs. The full-length (FL) protein consists of an N-terminal (blue), C-terminal (orange) and central SELOR domain (purple). The construct 1-480 has a C-terminal deletion, whereas in the construct 110-480 both the N- and C-terminus are truncated. Both deletion constructs were also rendered EJC-binding deficient by mutating the amino acid residues 188 and 218 (F188D, W218D). **D**: Relative quantification of the CLN6 (top) and TOE1 (bottom) transcript isoforms by qPCR in the indicated cell lines. The V5-tagged rescue proteins expressed in the KO condition are shown schematically in Figure 6C. Rescue protein expression is confirmed in Figure 6E and F. Individual data points and means are plotted from n=3 experiments. **E and F:** Western blot of samples shown in Figure 6D. The expression of rescue proteins was confirmed by an antibody against CASC3 (E) and an antibody recognizing the V5 tag (F).

Finally, CASC3 deletion mutants were expressed in the CASC3 KO cells to identify the minimal part necessary to rescue the effects on endogenous NMD targets (Figure 6D-F). As in the tethering experiment, the expression of full-length CASC3 and the C-terminal truncated variant 1-480 resulted in transcript isoform levels comparable to wild-type cells for the targets CLN6 and TOE1 (Figure 6D lanes 1-4). An EJC binding-deficient mutant of CASC3 (188/218 double point mutation) was unable to rescue, supporting the notion that CASC3 is recruited to the mRNA by binding to the EJC (Figure 6D, lane 5). Deleting the N-terminal 109 amino acids of CASC3 (110-480) did not alter the rescue ability (Figure 6D, lane 6). While in the tethering assay it was sufficient to place the N-terminus downstream of a termination codon, this part of CASC3 was not necessary to rescue NMD activity in the KO cells. This suggests that different domains of CASC3 act in a redundant manner during the activation of NMD by the EJC.

## Discussion

The role of CASC3 within the EJC has been the subject of scientific controversy for many years. CASC3 has been initially described as an EJC core protein, because it was required for the assembly of the EJC from recombinant protein components *in vitro* (21). However, it has been demonstrated that the mechanism of EJC assembly using recombinant proteins is mechanistically different from EJC assembly in splicing extracts or in living cell (11). Furthermore, several recent publications challenged the view of CASC3 being an EJC core component. For instance, CASC3 was reported to be present in substoichiometric amounts compared to the other three EJC core proteins EIF4A3, RBM8A, and MAGOH in HEK293 (7) and U2OS cells (81). Also, during mouse embryonic brain development CASC3 deficiency results in a different phenotype than the other EJC core components (82). By using CASC3 CRISPR-Cas9 knockout cells, we unambiguously establish that CASC3 is not required for EJC assembly or EJC-regulated splicing in the nucleus (Figure 7). Even previously reported CASC3-dependent alternative splice events (71) were not detectable in any of our KO or KD conditions. Therefore, our molecular analyses fully support the recently emerging view of defining CASC3 as a peripheral EJC component. As a mainly cytoplasmic component of the EJC we propose that the principal role of CASC3 is to alter the efficiency by which NMD-sensitive transcript isoforms are degraded.

**Figure 7.**
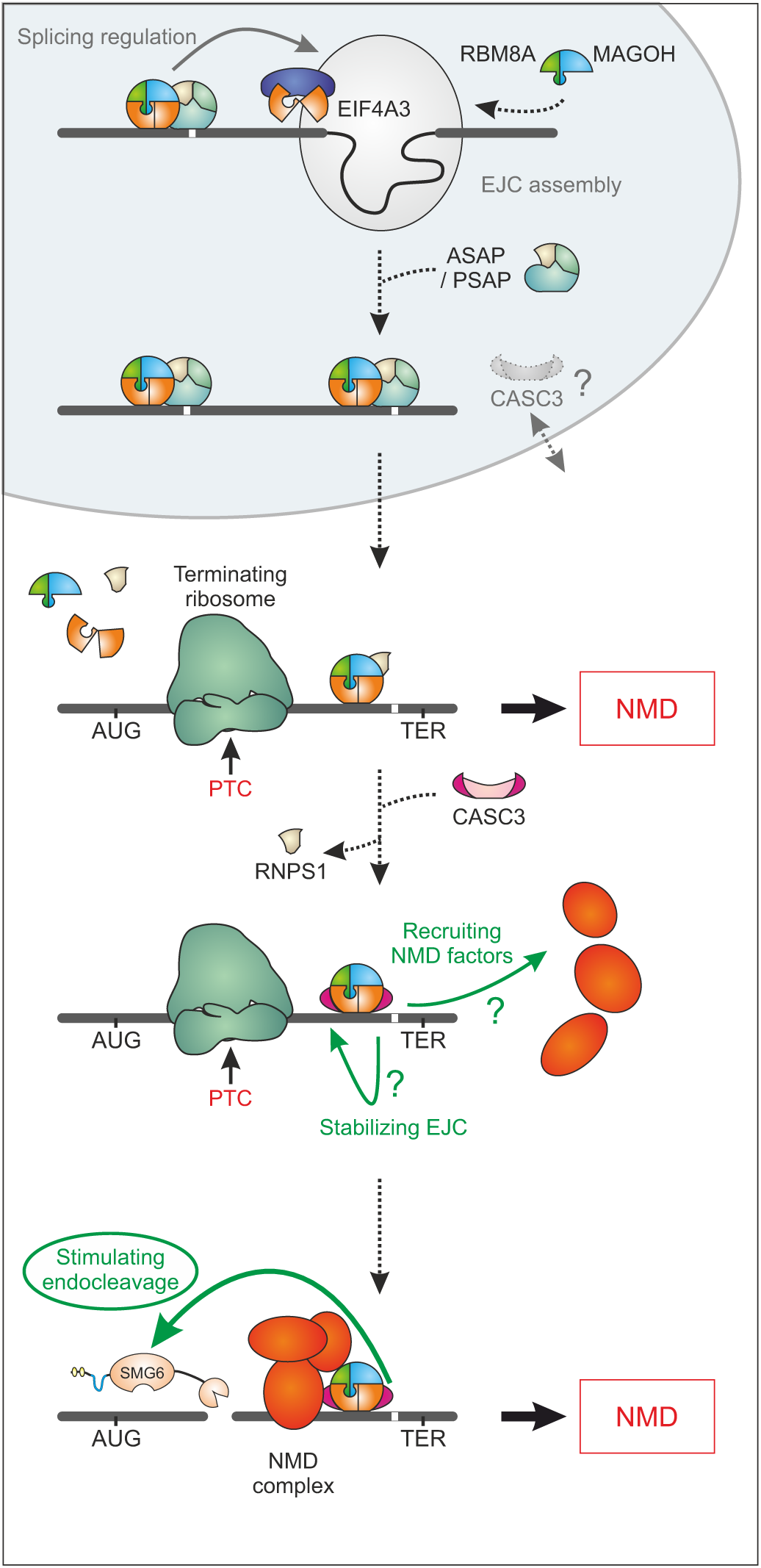
Model of CASC3’s cellular function. Schematic depiction of the proposed function of CASC3 in the cell. We have found no evidence that CASC3 is necessary for EJC assembly in the nucleus, although CASC3 shuttles between cytoplasm and nucleus. Transcripts that require the deposition of the EJC to be correctly spliced were not affected by a lack of CASC3. In the cytoplasm, premature termination codon (PTC)- containing transcripts may still be degraded by NMD during the initial round(s) of translation in a CASC3-independent manner. CASC3 association with the EJC maintains and/or promotes the NMD-stimulating effect of the EJC, resulting in the degradation of transcripts that evaded initial NMD activation.

Although NMD has been extensively studied in the past decades and many NMD factors have been identified and characterized, no universal model exists that describes how they work together to elicit NMD. While a function of CASC3 in NMD has been reported before, previous analyses did not show consistent results, ranging from a substantial contribution of CASC3 to only a minor role in NMD (12,17,22,34). In addition, none of the previous publications performed transcriptome-wide analyses but concentrated on reporter mRNAs of only a few selected endogenous NMD targets. We reasoned that the inconsistent results in the literature may be influenced by variable CASC3 knockdown efficiency. By generating CASC3 knockout cell lines, we can for the first time analyze the global effects of a complete depletion of CASC3 on the transcriptome and can exclude that residual CASC3 protein masks these effects. Interestingly, the CASC3 knockout cells appear phenotypically normal, unlike when the EJC components EIF4A3, RBM8A, or MAGOH are depleted. Nonetheless, since CASC3 is required for mouse embryogenesis and involved in the transport of mRNAs in *D. melanogaster*, it is likely that CASC3 downregulation in highly specialized cell types such as neurons or in developing tissues would result in a more severe phenotype compared to HEK293 cells.

In recent years, high-throughput RNA-sequencing became an increasingly important method for the analysis of NMD. Several RNA-Seq datasets of cells with NMD-factor knockdowns have been generated and analyzed (31,33,83–85). However, these datasets were obtained in different cell lines, with different amounts of replicates and due to the rapid developments of next-generation sequencing, not using the same technologies. Furthermore, batch effects and divergent approaches of data analyses may contribute to the fact that only a minor overlap of NMD targets could be established so far (31). We compared the results of our CASC3 KO RNA-sequencings to the most recent and comprehensive NMD factor analysis performed by Colombo *et al.* (31). The differential expression analysis revealed that many of the top upregulated genes in the CASC3 KO datasets are also significantly affected by UPF1 or SMG6/7 knockdowns (31) and/or encode for small RNA (sRNA) host genes, a previously described class of NMD targets (33).

We detected many alternative splicing events in the CASC3 KO data, which was unexpected given that CASC3 was apparently dispensable for the nuclear EJC-related functions. However, we could attribute these splicing patterns to dysfunctional NMD, since isoform-specific algorithms revealed that predominantly PTC-containing transcripts accumulated. A comprehensive bioinformatics analysis workflow and a systematic approach to detect affected transcripts under NMD factor knockdown/knockout conditions could therefore be a crucial step to paint a complete picture of the regulation of transcripts by NMD in the future. Our initial screen of the KO cells showed that compared to the siRNA-mediated CASC3 KD in WT cells, the genomic CASC3 KO cells demonstrated a more pronounced NMD inhibition. It is therefore important to reduce the amount of residual CASC3 protein as much as possible to obtain consistent and robust effects.

How exactly CASC3 activates NMD when bound to an EJC, is not yet fully understood. Previously, we reported that the presence of EJCs in the 3′ UTR enhances endonucleolytic cleavage (32). In line with the proposed role as a peripheral NMD-activating EJC component, we observed that CASC3 stimulates SMG6-dependent endonucleolytic cleavage, thereby promoting the degradation of NMD-targeted transcripts. This effect can be recapitulated by tethering full-length CASC3, its N-terminal two thirds (1-480) or just its N-terminal 137 amino acids to a reporter mRNA. How the small N-terminal region, which cannot assemble into the EJC or contains any known protein domains or sequence motifs can elicit NMD remains to be determined. It is also unclear, if the N-terminus activates translation-dependent degradation, as it was previously shown for the full length CASC3 (79). Since the N-terminal segment of CASC3 is a region of low-complexity it could hypothetically undergo liquid-liquid phase separation (LLPS) and be present in condensates with mRNA decay factors, such as processing bodies (P-bodies). In agreement with this idea CASC3 was shown to localize to cytoplasmic granules when overexpressed (86).

Our data suggest that CASC3 activates NMD by potentially redundant mechanisms. Binding of CASC3 to the EJC could have an indirect effect on NMD stimulation by increasing the stability of the bound EJC and thus maintaining the possibility of efficient endonucleolytic cleavage of the transcript. An indication for this role comes from the initial *in vitro* observation that CASC3 stabilizes recombinant EJCs (21). Additionally, the moderately reduced pull-down of UPF3B with EIF4A3 could indicate that cytoplasmic NMD-competent EJCs are less stable in CASC3 KO cells. Alternatively, CASC3 may directly contribute to the recruitment of NMD factors. We therefore propose that CASC3 potentially in conjunction with UPF3B links the EJC with the NMD machinery. In particular, CASC3 influences the contribution of SMG6-mediated endonucleolytic and SMG7-dependent exonucleolytic decay pathways to the overall degradation efficiency of NMD. Accordingly, in wild type cells a knockdown of SMG7 only had a marginal effect on the abundance of the analyzed NMD reporter mRNA, whereas it clearly impaired NMD in CASC3 KO cells. Also, the amount of endonucleolytic cleavage-derived 3′ fragments was reduced when CASC3 is depleted, mirroring the SMG6-knockdown condition.

By integrating CASC3 as a specific NMD-activating factor we can now postulate a modified model of EJC-dependent NMD, which is also compatible with several molecular properties of the EJC (Figure 7). Since CASC3 is only present in modest amounts in the cytoplasm, it will probably not immediately associate with all EJCs on recently exported mRNPs. This would also not be necessary, since most EJCs are located in the coding sequence and will therefore be removed by the first translating ribosome. However, mRNAs containing PTCs will carry one or more EJCs in their 3′ UTR, which are available for binding of CASC3. The first few translating ribosomes may terminate upstream of CASC3-free EJCs, which could preferentially trigger SMG7-dependent exonucleolytic degradation. Previously, NMD has been proposed to occur primarily in the pioneering round of translation when newly synthesized transcripts are bound to the cap-binding complex (87, 88). This model has been challenged and there is evidence that NMD can occur on already translating mRNAs and possibly with a constant probability during every round of translation (89–91). Thus, CASC3 could bind to the EJC at a later time point and then increase the probability to activate SMG6-mediated endonucleolytic degradation after each round of termination. Important molecular targets of CASC3 may be mRNAs that escape initial NMD activation, despite containing a PTC (89, 92). CASC3 may help to reduce the amount of these mRNAs by maintaining the NMD-activating function of the EJC, either by increasing its stability on the mRNA or via direct interactions with the NMD machinery. This concept would be consistent with the recent observation that NMD targets undergo several rounds of translation before endonucleolytic cleavage occurs (89).

In summary, our data paint a picture, in which CASC3 has no essential EJC-related function in the nucleus, but helps to sustain the EJC’s ability to induce NMD. We do not exclude the possibility that CASC3 is already associated with the EJC in the nucleus. However, our model of delayed binding of CASC3 to the EJC in the cytoplasm would explain why only a small amount of CASC3 is sufficient to activate EJC-dependent NMD. In this model, CASC3 is an indispensable cytoplasmic component of the EJC that helps to degrade mRNAs that failed to unload all their bound EJCs during the initial rounds of translation. Thus, the binding of CASC3 to the EJC could signal the final round(s) of translation of an mRNA.

## Data Availability

The datasets produced in this study are available in the following databases. These data will be made publicly accessible upon publication.

- RNA-seq data have been deposited in the ArrayExpress database (93) at EMBL-EBI under accession number E-MTAB-8461 (https://www.ebi.ac.uk/arrayexpress/experiments/E-MTAB-8461).
- The mass spectrometry proteomics data have been deposited to the ProteomeXchange Consortium via the PRIDE (94) partner repository with the dataset identifier PXD015754 (https://www.ebi.ac.uk/pride/archive/projects/PXD015754).

## Author Contributions

Conceptualization, N.H.G., J.V.G. and V.B.;

Methodology, N.H.G., V.B., J.V.G., and C.K.F.;

Software, T.B.B., V.B., J.L.W., S.K. and C.D.;

Investigation, J.V.G, V.B., J.L.W., S.K., D.U.A. and S.C.;

Resources and Data Curation, T.B.B., J.L.W., S.K., J.A. and C.D.;

Writing – Original Draft, Review & Editing, N.H.G., J.V.G. and V.B.;

Visualization, J.V.G., V.B. and T.B.B.;

Supervision, N.H.G., C.D. and M.K.;

Funding Acquisition, N.H.G. and C.D.

## Acknowledgements

We thank members of the Gehring lab for discussions and reading of the manuscript. We also thank Marek Franitza and Christian Becker (Cologne Center for Genomics, CCG) for preparing the sequencing libraries and operating the sequencer. We acknowledge Tobias Jakobi for helping with infrastructure support.

