## Supplementary Figures for "CASC3 promotes transcriptome-wide activation of nonsense-mediated decay by the exon junction complex"

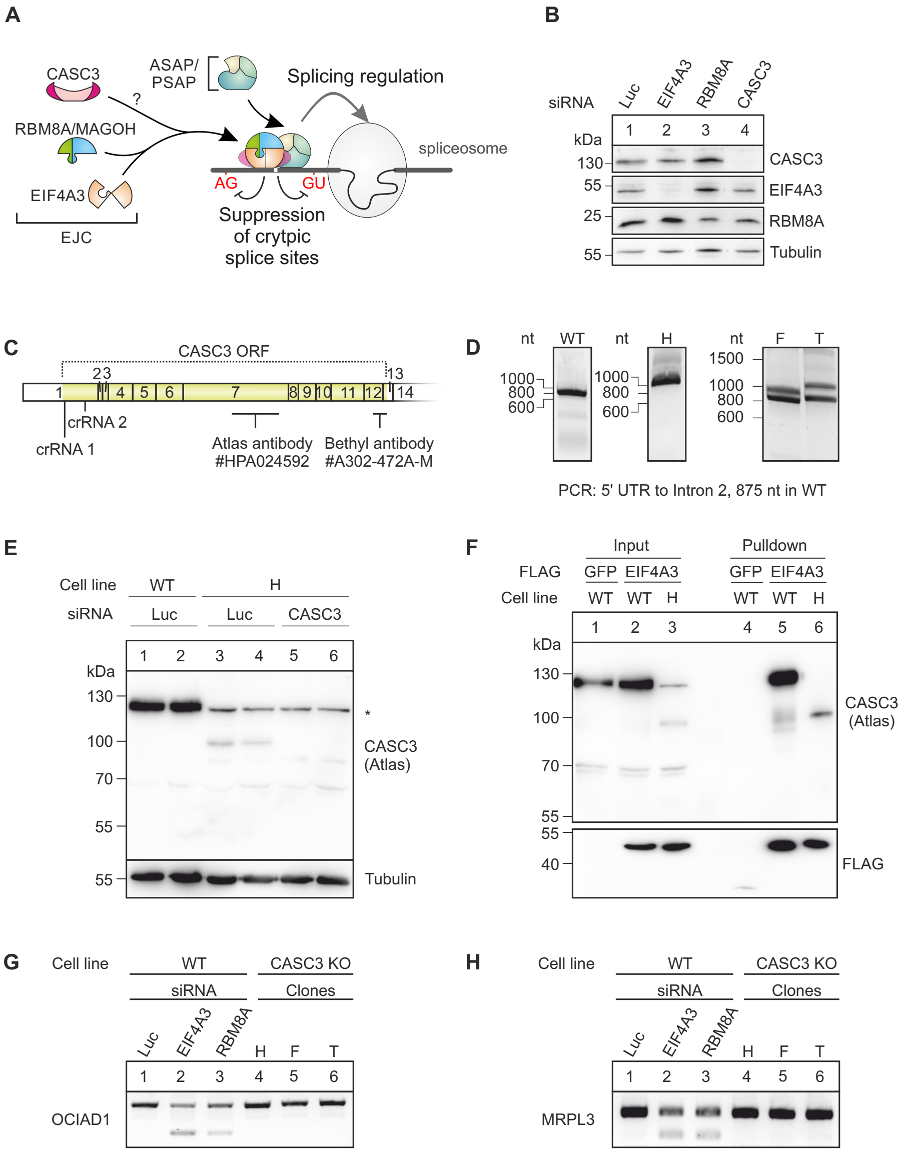


**Supplementary Figure 1 - Establishment and validation of CASC3 KO cell lines.**

**A:** Model of splicing regulation by the EJC, modified from Boehm *et al.* (2018).

**B:** Confirmation of the knockdowns shown in Figure 1A-D by western blotting.

**C**: Recognition sites of the two antibodies used in this study to detect CASC3. The coding exons 1-13 and a part of the non-coding exon 14 of CASC3 are depicted. The open reading frame (ORF) is marked yellow. The positions of the crRNA sequences to generate the knockout cells are also indicated.

**D**: Amplification of the genomic CASC3 locus targeted by CRISPR-Cas9 editing in the cell lines H, F and T. The sequences of the insertions were analyzed by cloning and Sanger-sequencing of the respective PCR-fragments and are shown in Figure 1F.

**E:** Western blot of cell lysates from WT cells and the CASC3 KO cell line H that have been treated with the indicated siRNAs.

**F:** Co-immunoprecipitation of FLAG-GFP and FLAG-EIF4A3 that were stably expressed in the indicated cell lines. Endogenous CASC3 and the recombinant FLAG proteins are detected by western blotting.

**G and H**: RT-PCR of transcript isoforms of the genes OCIAD1 (G) and MRPL3 (H) after siRNA-mediated knockdown of the indicated EJC components, Luciferase (Luc) as a negative control and in the CASC3 KO cell lines H, F and T.

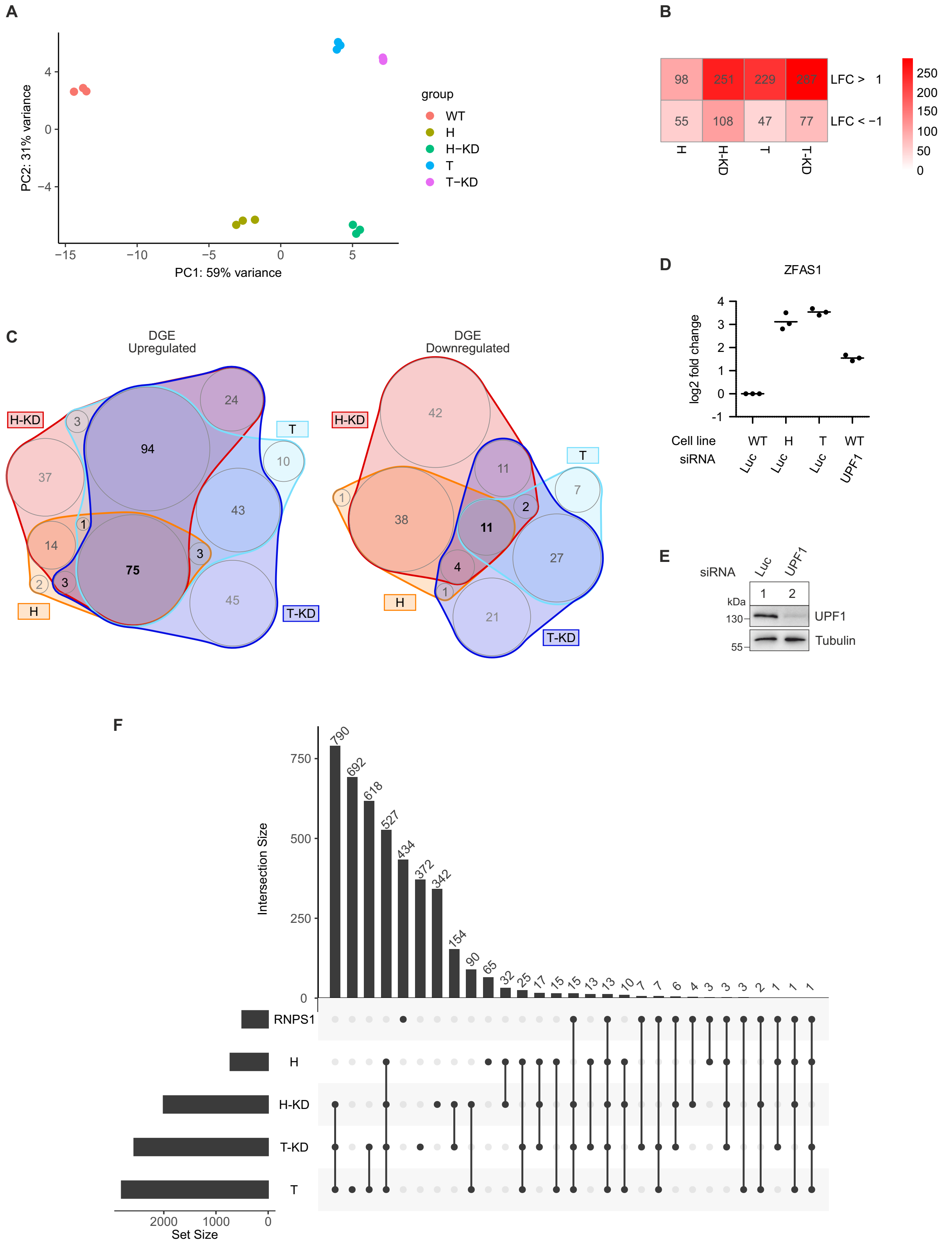


**Supplementary Figure 2 - Transcriptome-wide effects of the CASC3 KO.**

**A**: Principal component analysis (PCA) of differential gene expression data.

**B**: Summary of significantly up- or downregulated genes in the differential gene expression analysis with an absolute log2 fold change (|LFC|) > 1.

**C**: Overlap of all intersections of differentially expressed genes in all CASC3 KO conditions shown as nVenn plot.

**D**: Relative quantification by qPCR of the ZFAS1 transcript in WT cells, CASC3 KO cell lines H and T and WT cells treated with siRNA targeting UPF1. Individual data points and means are plotted (n=3).

**E**: Confirmation of the UPF1 knockdown by western blotting.

**F**: Upset plot of intersections of alternatively spliced junctions in the shown conditions.

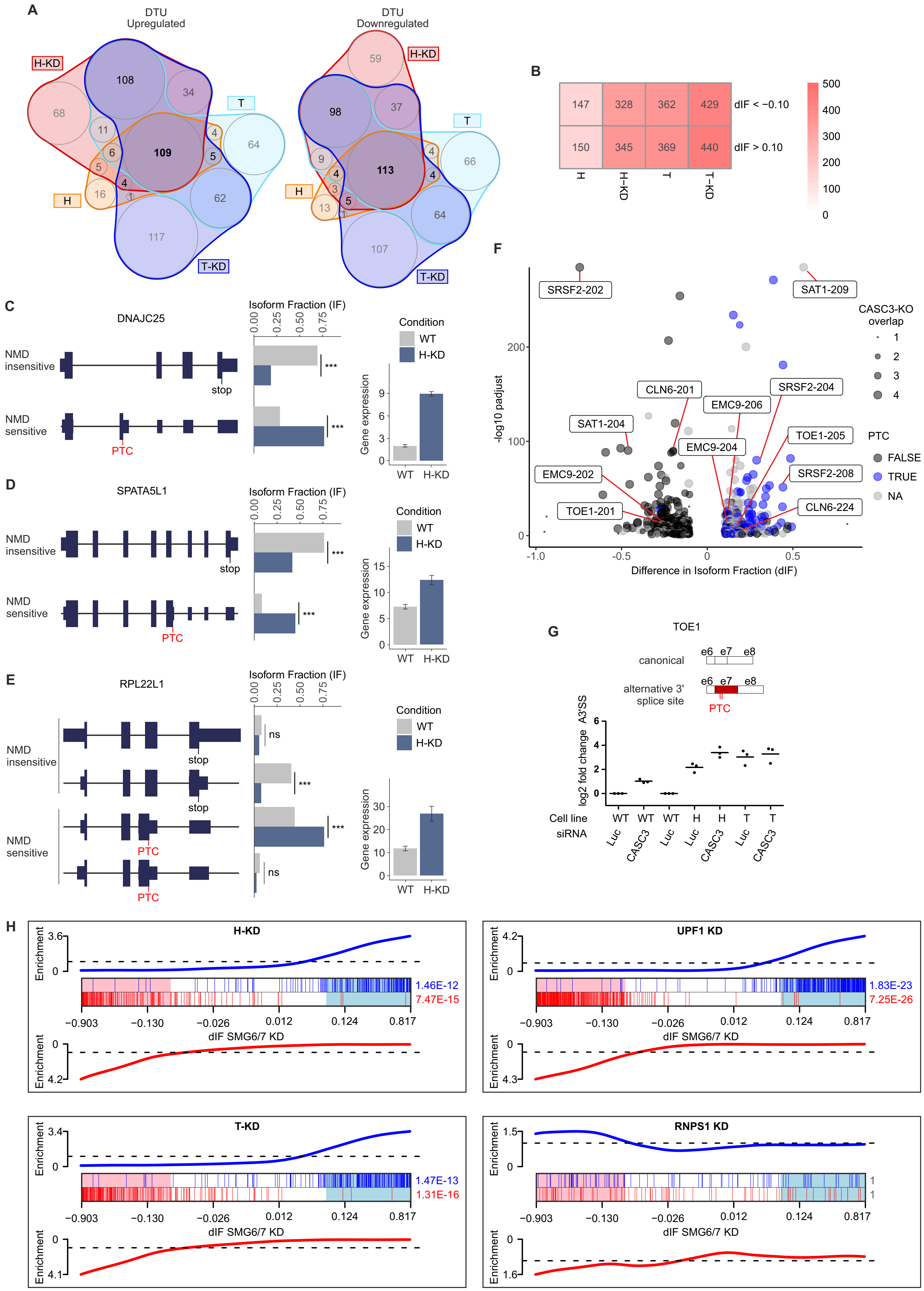
A
DTU Upregulated
DTU Downregulated
H-KD
T
H
T-KD
B
147
328
362
429
150
345
369
440
dIF < −0.10
dIF > 0.10
H
H-KD
T
T-KD
C
DNAJC25
NMD insensitive
NMD sensitive
stop
PTC
Isoform Fraction (IF)
Condition
WT
H-KD
Gene expression
D
SPATA5L1
E
RPL22L1
F
SRSF2-202
SAT1-209
CASC3-KO overlap
PTC
FALSE
TRUE
NA
-log10 padjust
Difference in Isoform Fraction (dIF)
G
TOE1
canonical
alternative 3' splice site
log2 fold change A3'SS
Cell line
siRNA
H
H-KD
UPF1 KD
T-KD
RNPS1 KD
Enrichment
dIF SMG6/7 KD
1.46E-12
7.47E-15
1.83E-23
7.25E-26
1.47E-13
1.31E-16

**Supplementary Figure 3 - Analysis of differentially expressed transcript isoforms in the CASC3 KO.**

**A**: Overlap of all intersections of transcript isoforms up-or downregulated in IsoformSwitch analysis shown as nVenn plot. DTU: Differential transcript usage.

**B**: Summary of transcript isoforms up- or downregulated in IsoformSwitch analysis with an absolute Difference in Isoform Fraction (|dIF|) > 0.1.

**C-E**: Quantification of transcript isoforms from DNAJC25 (C), SPATA5L1 (D), RPL22L1 (E) by IsoformSwitchAnalyzeR.

**F**: Volcano plot of IsoformSwitch data for the condition H-KD using overlap from Supplementary Figure S3A as point size definition and PTC status as color definition. Isoforms with no annotated open reading frame are designated as “NA”. Ensembl transcript names are indicated for the genes validated in Figure 3C-E and Supplementary Figure S3G. Difference in Isoform Fraction (dIF) is plotted against –log10 padjust (adjusted p-value).

**G**: Relative quantification of the schematically depicted transcript isoforms from the gene TOE1 by qPCR in WT cells treated with Luc siRNA as a negative control, WT cells treated with CASC3 siRNAs, CASC3 KO cell line H treated with Luc siRNA, CASC3 KO cell line H treated with CASC3 siRNAs and CASC3 KO cell line T treated with Luc siRNA as well as CASC3 KO cell line T treated with CASC3 siRNAs. Individual data points and means are plotted (n=3).

**H**: Barcode plots as shown in Figure 3G. The worms represent the relative enrichment of vertical bars in the plot.

Significant in WT/CTL, KO/CTL and KO/WT

| | WT/CTL | KO/CTL | KO/WT |
|---|---|---|---|
| CWC22 | 3.92 | 4.48 | 0.54 |
| ZC3H11A | 3.38 | 3.00 | -0.44 |
| SDE2 | 3.30 | 4.02 | 0.72 |
| CHTOP | 3.21 | 2.63 | -0.52 |
| DDX39B | 3.18 | 2.80 | -0.42 |
| PPWD1 | 2.72 | 3.53 | 0.69 |
| EIF4G1 | 1.96 | 1.21 | -0.71 |
| CCDC12 | 1.78 | 2.20 | 0.42 |
| DDX17 | 1.25 | 0.95 | -0.33 |

Significant in WT/CTL and KO/WT

| | | | |
|---|---|---|---|
| CASC3 | 4.74 | 0.29 | -4.34 |
| UPF3B | 3.97 | 2.45 | -1.52 |

Significant in KO/CTL and KO/WT

| | | | |
|---|---|---|---|
| USP15 | 0.10 | 0.28 | 0.14 |
| CMBL | 0.15 | -0.47 | -0.62 |
| WARS | 0.12 | -0.97 | -1.08 |

Significant in WT/CTL and KO/CTL

| | | | |
|---|---|---|---|
| EIF4A3 | 6.12 | 5.50 | -0.62 |
| PYCR2 | 5.00 | 3.47 | -1.57 |
| RBM8A | 4.92 | 4.86 | 0.02 |
| MAGOHB | 4.82 | 4.82 | 0.03 |
| ACIN1 | 4.60 | 3.78 | 0.07 |
| PNN | 4.49 | 4.50 | -0.01 |
| RNPS1 | 4.08 | 3.85 | 0.13 |
| CRNKL1 | 4.05 | 4.57 | 0.53 |
| THOC5 | 4.00 | 3.59 | -0.51 |
| ZC3H14 | 3.93 | 3.72 | -0.16 |
| SAP18 | 3.92 | 4.00 | 0.12 |
| KIF1C | 3.91 | 4.25 | 0.26 |
| DDX3X | 3.88 | 2.65 | -1.25 |
| EIF2A | 3.85 | 2.78 | -1.05 |
| CWC27 | 3.83 | 4.87 | 0.45 |
| SRRM2 | 3.78 | 3.56 | -0.37 |
| CCDC9 | 3.74 | 3.58 | -0.06 |
| CDC5L | 3.68 | 3.95 | 0.25 |
| THOC2 | 3.64 | 3.09 | -0.44 |
| PRPF19 | 3.55 | 3.79 | 0.25 |
| TRA2B | 3.53 | 3.44 | -0.12 |
| PLRG1 | 3.49 | 3.71 | 0.28 |
| PRPF4B | 3.49 | 3.85 | 0.43 |
| PDCD4 | 3.41 | 3.07 | -0.27 |
| CDC40 | 3.40 | 3.87 | 0.59 |
| PRPF38A | 3.34 | 3.18 | -0.16 |
| SNW1 | 3.33 | 3.51 | 0.27 |
| ARGLU1 | 3.33 | 3.22 | -0.11 |
| SRSF7 | 3.32 | 3.54 | 0.18 |
| SRSF6 | 3.29 | 3.26 | -0.03 |
| SNRNP40 | 3.29 | 3.50 | 0.23 |
| PAXBP1 | 3.27 | 3.32 | 0.14 |
| SNRNP200 | 3.27 | 3.51 | 0.21 |
| BUD13 | 3.27 | 3.49 | 0.37 |
| PRPF8 | 3.27 | 3.51 | 0.26 |
| DDX23 | 3.22 | 3.31 | 0.11 |
| EFTUD2 | 3.15 | 3.34 | 0.23 |

Significant in WT/CTL and KO/CTL continued

| | | | |
|---|---|---|---|
| TFIP11 | 3.12 | 3.16 | 0.10 |
| SART1 | 3.11 | 3.21 | 0.12 |
| XAB2 | 3.11 | 3.61 | 0.54 |
| CFAP20 | 3.08 | 2.74 | -0.39 |
| PABPN1 | 3.06 | 2.75 | -0.17 |
| RBM22 | 3.05 | 3.42 | 0.24 |
| MFAP1 | 3.00 | 3.08 | 0.06 |
| CD2BP2 | 2.99 | 2.93 | 0.09 |
| USP39 | 2.96 | 3.10 | 0.09 |
| RBM25 | 2.96 | 2.69 | -0.13 |
| SRSF1 | 2.94 | 2.90 | 0.01 |
| PRPF3 | 2.91 | 3.16 | 0.14 |
| PRPF6 | 2.90 | 3.06 | 0.23 |
| THOC1 | 2.90 | 2.63 | -0.29 |
| HIST1H2BN | 2.84 | 2.84 | 0.35 |
| IK | 2.83 | 2.95 | 0.09 |
| PRPF40A | 2.81 | 2.59 | 0.06 |
| HIST2H3PS2 | 2.77 | 2.65 | -0.25 |
| HIST1H1C | 2.74 | 2.53 | -0.33 |
| SMU1 | 2.73 | 2.67 | 0.02 |
| PRPF4 | 2.73 | 2.84 | 0.16 |
| SNRPA1 | 2.72 | 3.00 | 0.24 |
| SF3B2 | 2.66 | 2.83 | 0.13 |
| CTNNBL1 | 2.65 | 2.76 | 0.07 |
| SF3B3 | 2.52 | 2.70 | 0.10 |
| SF3B1 | 2.51 | 2.66 | 0.10 |
| TCERG1 | 2.51 | 2.14 | -0.39 |
| SF3A1 | 2.48 | 2.64 | 0.14 |
| WBP11 | 2.46 | 2.37 | -0.15 |
| FNBP4 | 2.45 | 2.73 | 0.31 |
| DHX15 | 2.38 | 2.18 | -0.14 |
| SNRPB2 | 2.31 | 2.49 | 0.20 |
| SF3A3 | 2.28 | 2.46 | 0.18 |
| RBMX | 2.18 | 1.63 | -0.56 |
| SNRPD2 | 2.15 | 2.31 | 0.15 |
| SF3A2 | 2.08 | 2.32 | 0.14 |
| CHERP | 1.99 | 2.03 | 0.03 |
| SNRPD1 | 1.90 | 2.04 | 0.15 |
| U2SURP | 1.90 | 1.72 | -0.13 |
| BCLAF1 | 1.90 | 2.19 | 0.25 |
| LUC7L3 | 1.86 | 1.94 | 0.26 |
| HNRNPM | 1.79 | 1.59 | -0.17 |
| XRCC6 | 1.76 | 1.26 | -0.33 |
| XRCC5 | 1.74 | 1.36 | -0.31 |
| THRAP3 | 1.60 | 2.25 | 0.65 |
| DHX38 | 1.59 | 1.74 | 0.11 |
| SRPK1 | 1.58 | 1.36 | -0.18 |
| PRKDC | 1.56 | 1.55 | 0.07 |
| SRRT | 1.56 | 1.41 | -0.22 |
| SARNP | 1.28 | 1.09 | -0.34 |
| EIF3G | 1.20 | 0.79 | -0.51 |
| SAFB | 1.18 | 0.89 | -0.32 |
| TAF15 | 1.01 | 0.91 | -0.09 |
| DDX5 | 0.96 | 0.89 | -0.03 |
| SNRPD3 | 0.70 | 0.98 | 0.26 |
| USP9X | 0.53 | 0.72 | 0.01 |

Significant in WT/CTL only

| | | | |
|---|---|---|---|
| PYCR1 | 4.45 | 2.29 | -1.83 |
| PPIG | 3.51 | 3.38 | 0.21 |
| HIST1H4A | 2.73 | 2.69 | -0.06 |
| RPL8 | 2.41 | 1.86 | -0.66 |
| DHX8 | 2.28 | 2.57 | 0.60 |
| CDK11B | 2.24 | 2.82 | 0.57 |
| RPL7A | 2.20 | 1.40 | -0.76 |
| RPL13 | 2.11 | 1.48 | -0.77 |
| RPL6 | 1.99 | 1.18 | -0.91 |
| RPL7 | 1.87 | 0.82 | -0.71 |
| PARP1 | 1.86 | 2.03 | 0.08 |
| EIF3A | 1.82 | 0.88 | -0.96 |
| EIF3C | 1.81 | 0.93 | -0.98 |
| RPS26 | 1.77 | 0.55 | -1.23 |
| RBM17 | 1.77 | 1.43 | -0.40 |
| CSNK2A1 | 1.63 | 1.08 | -0.40 |
| RPL34 | 1.56 | 0.83 | -0.66 |
| RPS8 | 1.54 | 0.82 | -0.61 |
| RPS9 | 1.52 | 0.96 | -0.63 |
| RPS14 | 1.51 | 0.85 | -0.74 |
| RPS4X | 1.46 | 0.77 | -0.61 |
| RPS7 | 1.45 | 0.78 | -0.57 |
| RPS3A | 1.43 | 0.76 | -0.68 |
| EIF3I | 1.39 | 0.67 | -0.87 |
| RPS20 | 1.29 | 0.70 | -0.55 |
| RPS25 | 1.29 | 0.69 | -0.63 |
| RPS3 | 1.28 | 0.66 | -0.66 |
| RPS16 | 1.28 | 0.65 | -0.66 |
| RPSA | 1.26 | 0.62 | -0.68 |
| U2AF2 | 1.26 | 1.39 | -0.04 |
| RPS18 | 1.24 | 0.53 | -0.75 |
| GNB2L1 | 1.20 | 0.60 | -0.65 |
| LUC7L2 | 1.19 | 1.00 | -0.10 |
| SNRNP70 | 1.01 | 0.66 | -0.36 |
| KHDRBS1 | 0.89 | 0.25 | -0.68 |
| IRS4 | 0.75 | -0.44 | -1.19 |
| UBB | 0.62 | 0.25 | -0.44 |
| FUS | 0.44 | 0.74 | 0.30 |
| NONO | 0.44 | -0.14 | -0.59 |
| SFPQ | 0.37 | -0.07 | -0.47 |
| NSRP1 | 0.36 | 0.74 | 0.43 |
| GEMIN5 | 0.21 | -0.07 | -0.27 |
| HNRNPA1 | 0.14 | -0.05 | -0.22 |
| ANKFY1 | -0.32 | 0.21 | 0.49 |

Significant in KO/CTL only

| | | | |
|---|---|---|---|
| AQR | 4.85 | 4.10 | 0.55 |
| SON | 3.14 | 3.18 | 0.02 |
| RSRC1 | 3.45 | 3.04 | -0.12 |
| SFSWAP | 3.20 | 2.87 | -0.04 |
| RBM39 | 2.11 | 2.25 | -0.02 |
| FAT3 | 1.69 | 1.78 | 0.00 |
| SNRPN | 0.45 | 0.92 | 0.35 |
| HSPA8 | 0.44 | 0.34 | -0.07 |
| HSPH1 | 0.34 | 0.33 | -0.14 |
| HSPA1B | 0.69 | 0.31 | -0.38 |
| SMC1A | 0.28 | 0.17 | -0.12 |
| HDAC6 | 0.69 | -0.56 | -0.72 |

**Supplementary Figure 4 - Overview of mass spectrometry results.** All targets are listed that are shown in the volcano plots in Figure 4. The log2 fold changes of the WT/CTL (first column), KO/CTL (second column) and KO/WT (third column) comparisons are shown. The color of points corresponds to the volcano plots in Figure 4. Full results are shown in Supplementary Table S2.

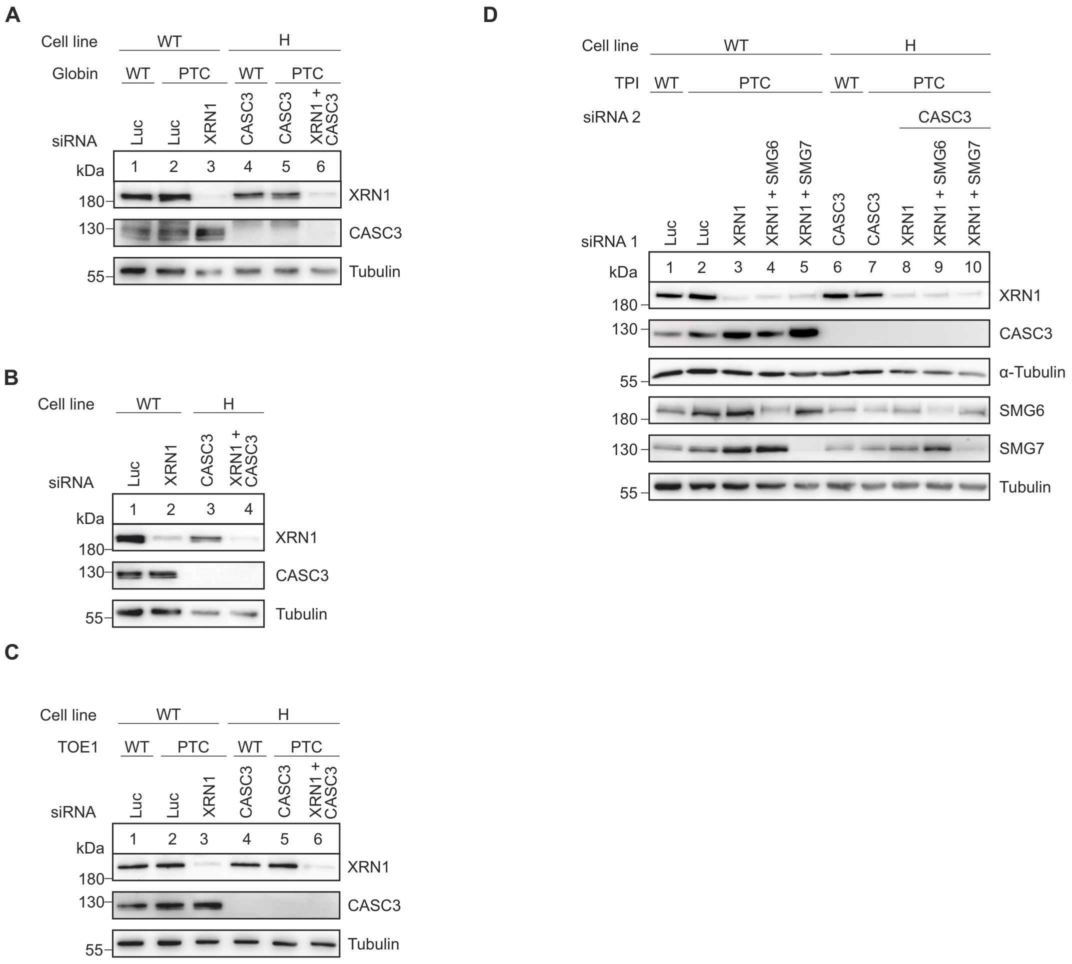


**Supplementary Figure 5 - Confirmation of knockdowns performed for NMD reporter assays.**

**A**: Western blot of samples shown in Figure 5B.

**B**: Western blot of samples shown in Figure 5D.

**C**: Western blot of samples shown in Figure 5E.

**D**: Western blot of samples shown in Figure 5G.

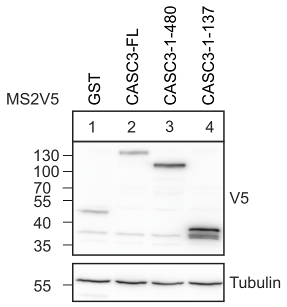


**Supplementary Figure 6 - Confirmation of tethering protein expression.** Western blot of samples shown in Figure 6B.
